# Histone Variant macroH2A1 Spatially and Functionally Organizes Human Papillomavirus Replication Foci in the Productive Stage of Infection

**DOI:** 10.1101/2021.07.27.453825

**Authors:** Simran Khurana, Tovah E. Markowitz, Juraj Kabat, Alison A. McBride

## Abstract

The life cycle of HPV depends on keratinocyte differentiation as the virus modulates and takes advantage of cellular pathways to replicate its genome and assemble viral particles in differentiated cells. Viral genomes are amplified in nuclear replication foci in differentiated keratinocytes, and DNA repair factors from the DNA damage response signaling pathway are recruited to replicate viral DNA. The HPV genome is associated with cellular histones at all stages of the infectious cycle, and here we show the histone variant macroH2A1 is bound to the HPV genome and enriched in viral replication foci in differentiated cells. MacroH2A1 isoforms play important roles in cellular transcriptional repression, double strand break repair, and replication stress. The viral E8^E2 protein also binds to the HPV genome and inhibits viral replication and gene expression by recruiting NCoR/SMRT complexes. We show here that E8^E2 and SMRT also localize within replication foci, though not through direct interaction with macroH2A1. Conversely, transcription complexes containing RNA polymerase II and Brd4 are located on the surface of the foci. Foci generated with an HPV16 E8^E2 mutant genome are not enriched for SMRT or macroH2A1 but contain transcriptional complexes throughout the foci. We demonstrate that macroH2A1 promotes viral late transcription and propose that it does so by spatially separating replication and transcription activities within replication foci.

**IMPORTANCE:** Human papillomaviruses are small DNA viruses that cause chronic infection of cutaneous and mucosal epithelium. In some cases, persistent infection with HPV can result in cancer, and 5% human cancers are the result of HPV infection. In differentiated cells, HPV amplifies viral DNA in nuclear replication factories and transcribes late mRNAs to produce capsid proteins. However, very little is known about the spatial organization of these activities in the nucleus. Here we show that repressive viral and cellular factors localize within the foci to supress viral transcription, while active transcription takes place on the surface. The cellular histone variant macroH2A1 is important for this spatial organization.

## Introduction

High oncogenic risk human papillomaviruses (HR-HPVs), such as HPV16, 18, and 31, are the causative agents of anogenital and oropharyngeal cancers (1). In addition, infection with some HPV types from the Betapapillomavirus genus may predispose to squamous cell skin cancer (2). HPVs infect the actively proliferating basal layer of keratinocytes to establish a persistent infection (3). There are three stages of DNA replication in the HPV viral life cycle. First, there is an initial burst of viral DNA replication in the initial host cell and the viral genome becomes established as a low copy number extrachromosomal plasmid. During the second stage, established genomes are replicated and partitioned along with host DNA to daughter cells. Finally, when the infected cells differentiate, the viral genome amplifies to very high levels (4). During the productive phase of HPV viral life cycle, repair factors from the ATM and ATR (ataxia-telangiectasia mutated ATM and Rad3 related) DNA damage signaling pathways are hijacked by HPV to amplify viral DNA in non-dividing cells (5).

The viral E1 and E2 proteins initiate viral DNA replication and, in addition, E2 regulates transcription and facilitates partitioning of viral genomes (6). E1 is a helicase that unwinds the viral origin and recruits host cellular factors to the viral replication foci and co-expression of E1 and E2 proteins leads to the formation of nuclear foci that recruit DNA damage factors including pATM, pChk2, γH2AX, MRE11, and NBS1 (7–9). Additionally, the HPV E2 protein and cellular BRD4 proteins associate with and nucleate the formation of viral foci near common fragile sites of the host genome (10). Independently, the E7 protein activates the ATR and ATM signaling pathways both directly (5, 11, 12), and indirectly by inducing cellular proliferation that results in nucleotide deficiency and replication stress (13, 14). All papillomaviruses encode a fusion protein, E8^E2, that restricts viral genome replication and transcription (15, 16). HPV16 E8 mutant genomes over replicate in undifferentiated cells, and express increased levels of viral transcripts and late proteins in differentiated cells, as compared to the wild-type virus (15, 16). The E8^E2 protein competitively binds to E2BS (E2 binding sites) in the viral genome, and interacts with the host corepressor SMRT/NCoR complexes to regulate viral replication and transcription (16, 17).

The HPV genome is associated with cellular histones both in infected cells and in virion particles (18–20). Histones are post-translationally modified by acetylation, phosphorylation, methylation, sumoylation, and ubiquitination and these modifications affect chromatin accessibility and impact cellular and HPV gene expression (21, 22). In addition, variants of the canonical histones are associated with different cellular processes, but it is not known whether they also bind to the HPV genome and influence the viral life cycle. One such variant is macroH2A; macroH2A1 is a variant of the canonical H2A histone with a unique C-terminal 30kDa macro domain. The macroH2A1 encoding gene *H2AFY* encodes two splice variants, macroH2A1.1 and macroH2A1.2, which differ in only 30 amino acids in the carboxyl-terminal macro domain. This results in the formation of a poly-ADP-ribose (PAR) binding pocket in macroH2A1.1, but not in macroH2A1.2 (23). MacroH2A1.1 is predominantly expressed in differentiated cells while macroH2A1.2 is ubiquitously expressed in both differentiated and proliferating cells. (24, 25). Both macroH2A1.1 and macroH2A1.2 isoforms are recruited to sites of DNA damage and are involved in either non-homologous end joining (NHEJ) and/or homologous recombination (HR) (23, 26). MacroH2A1.2 also accumulates at common fragile sites upon replication stress (27).

Here, we examine the role of macroH2A1 on HPV genome amplification and transcription during the productive phase of the viral life cycle and show that macroH2A1 associates with HPV replication factories (foci). Depletion of macroH2A1 by siRNA did not affect viral replication but decreased levels of viral transcripts. We show that components of the cellular transcriptional machinery (including RNA Pol II Ser 5, RNA Pol II Ser 2 and Brd4) are present predominantly at the periphery of the replication foci. However, when macroH2A1 is absent from the replication foci, the cellular transcriptional machinery is localized to the interior of the foci, suggesting a role for macroH2A1 in the spatial separation of viral replication and transcription processes during the HPV productive viral life cycle.

## Results

### MacroH2A1 associates with HPV 18 and HPV31 viral replication foci

During a preliminary screen to determine whether histones with specific modifications were increased in the chromatin of HPV replication foci, we noted that the variant histone macroH2A1 was highly enriched at HPV31 replication foci in differentiated CIN-612 9E cells (designated 9E from here on). 9E cells are derived from a CINI cervical lesion and contain extrachromosomal HPV31 genomes that can amplify to high copy number in nuclear foci in differentiated cells (4). Both 9E cells and an HPV negative keratinocyte cell line (NIKS) were cultured on glass coverslips until confluent and differentiated in high calcium medium for five days. Differentiation of these cells results in amplification of viral DNA inside nuclear replication foci. As shown in Figure 1A and 1B, viral replication foci (identified by replication protein A, RPA, single strand DNA binding protein staining) were highly enriched for macroH2A1.2 in 9E cells compared to the rest of the nuclei. We observed macroH2A1.2 enrichment in ∼95% of replication foci and this was irrespective of their size. In contrast, macroH2A1.2 was generally diffuse in control NIKS cells.

**Figure 1.**
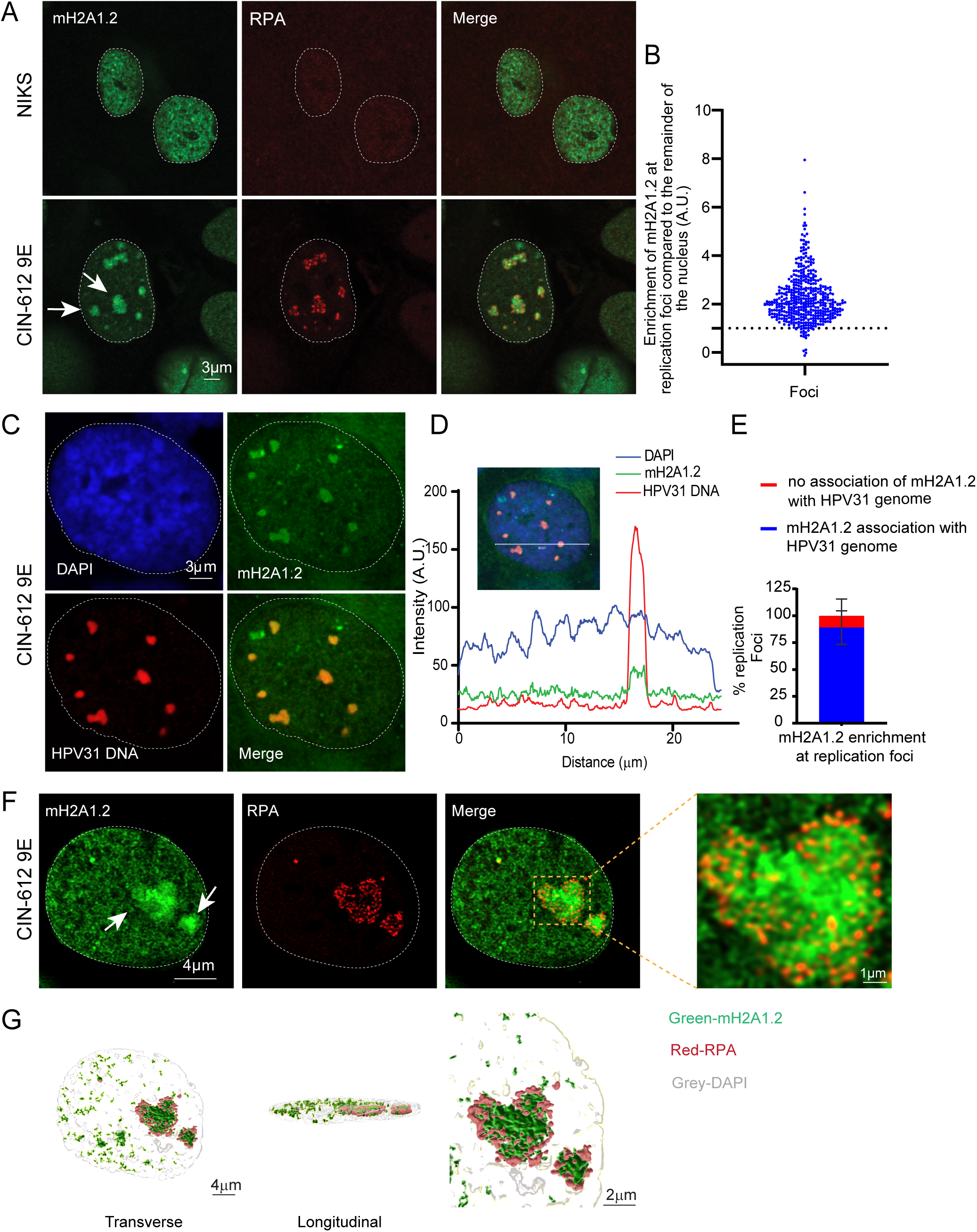
MacroH2A1.2 localizes to HPV31 replication foci in 9E cells. **A**. Differentiated NIKS or 9E cells were immunostained with antibodies against macroH2A1.2 (green) and RPA (red) and nuclei were counter stained with DAPI (outlined with dotted line). **B.** The mean fluorescent Intensity of macroH2A1.2 at the foci was calculated using ImageJ (values >1.0 indicate enrichment). In 9E cells (N=67), 424/443 RPA positive foci (∼95%) showed enrichment of macroH2A1.2, in three independent experiments. No RPA foci were observed In NIKS cells (N=213). White arrows indicate different sized foci. A.U.: Arbitrary unit. **C.** Differentiated 9E cells were analyzed by combined IF-FISH for colocalization of macroH2A1.2 (green) and HPV31 DNA (red). Nuclei were stained with DAPI. **D**. Fluorescence intensity line scan obtained by drawing a line through the nucleus shown in B using Leica LAS X software. **E**. The % of replication foci detected by HPV31 DNA FISH with macroH2A1.2 enrichment A total of 35 cells counted (N=35) and 172 HPV31 positive foci were scored in two independent experiments. **F**. High resolution image of deconvolved image is shown from single slice of Z stacks collected throughout the nucleus. White arrows indicate different sized foci. The magnified image in the box area demonstrates macroH2A1.2 association with RPA at viral foci. **G**. 3D reconstruction of RPA (red) and macroH2A1.2 (green) staining inside a replication factory. Surface-rendering was generated in IMARIS from a Z-stack image collected at optimum X, Y and Z settings and deconvolved in Huygens.

To verify that the RPA foci observed in 9E cells were in fact HPV replication factories, we confirmed the association of macroH2A1 with the viral genome using immunofluorescent staining for macroH2A1.2 followed by Fluorescence In situ Hybridization (FISH) for the HPV31 genome. Similar to the data shown in Figure 1A, we observed that ∼85% of HPV31 DNA positive replication foci colocalized with macroH2A1.2 (Figure 1C, D and E). High resolution confocal imaging and 3D image reconstruction showed that macroH2A1 localized in a diffuse pattern throughout most of the foci as compared to the punctate pattern of RPA (Figure 1F and 1G). We also examined macroH2A1.2 enrichment at replication factories in a NIKS cell line containing extrachromosomal HPV18 genomes and verified that ∼ 98% replication foci showed enrichment of macroH2A1.2 (Figure 2A and 2B) compared to the rest of the nuclei.

**Figure 2.**
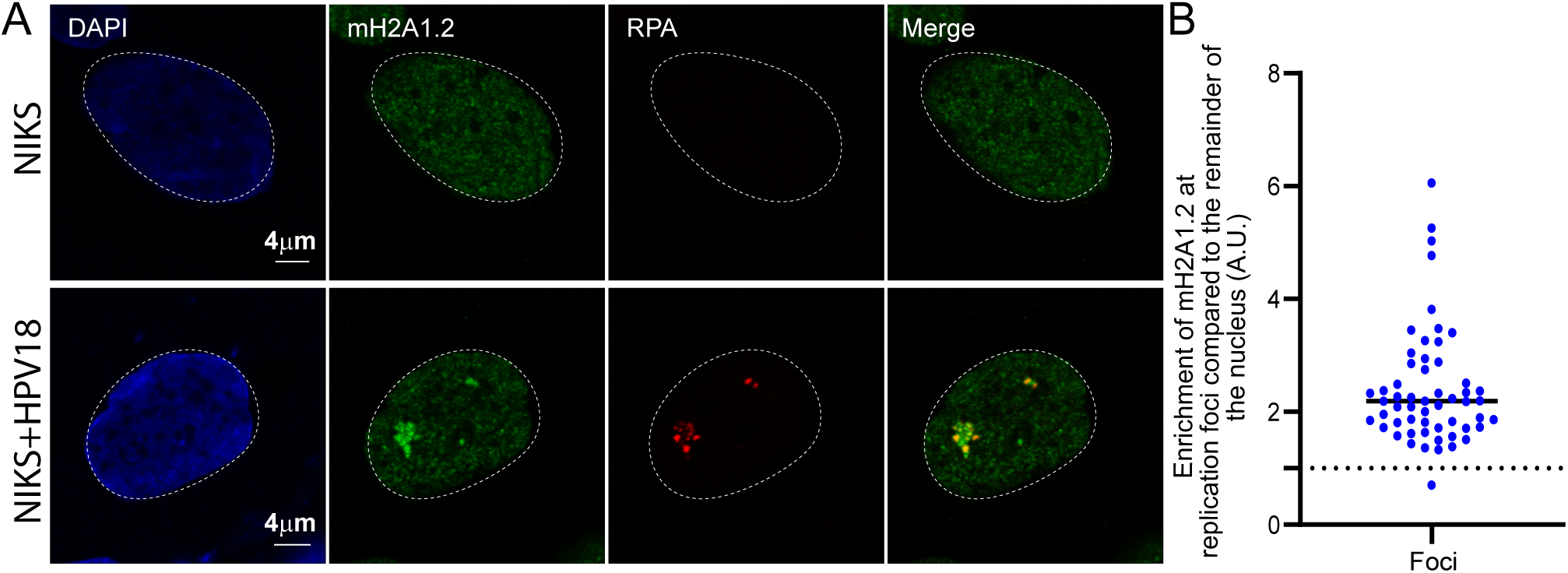
MacroH2A1.2 localizes to HPV18 replication foci. **A.** Differentiated NIKS, or NIKS containing extrachromosomal HPV18 genomes (NIKS-HPV18) were immunostained with antibodies against macroH2A1.2 (green) and RPA (red) and nuclei were stained with DAPI (blue) in two independent experiments. **B.** The mean fluorescent Intensity of macroH2A1.2 at the foci was calculated using ImageJ (values >1.0 indicate enrichment). In NIKS-HPV18 cells (N=41), 98% (55/56) RPA positive foci showed enrichment of macroH2A1.2. In NIKS, 0/65 cells contained RPA positive replication foci.

The splice variant macroH2A1.1 was also enriched at viral replication factories in 9E cells (Supplementary Figure 1A). In contrast, there was no enrichment of the core histones H2A, H3 or H4 in replication foci compared to the rest of the nucleus showing that this was not simply due to a general increased nucleosomal density (Supplementary Figure 1B and C). Our studies rely heavily on the specificity of the macroH2A1.1 and 1.2 antibodies. Therefore, the expression of macroH2A1.1. and 1.2 was downregulated with siRNA in 9E cells, followed by immunofluorescence and western blotting with the macroH2A1.1 and 1.2 antibodies (Supplementary Figure 2). This confirmed that the macroH2A1.1 and 1.2 antibodies were specific and did not cross react with viral or cellular proteins.

### The association of macroH2A1 at replication foci does not correlate with the presence of repressive histone modifications

MacroH2A1.2 is often associated with chromatin containing the repressive histone modifications H3K9me2/3 (26, 28). Therefore, we examined the localization of H3K9 dimethylated and H3K9 trimethylated histones in differentiated 9E cells and found that, unlike macroH2A1, these modifications were not enriched at viral replication foci (Supplementary Figure 3). Together, these findings indicate that the macroH2A1 proteins accumulate at viral replication foci, but this enrichment does not correlate with H3K9me2/3 associated repressive chromatin.

### Binding of macroH2A1.2 to viral chromatin increases in differentiated cells

Next, to determine whether macroH2A1.2 is incorporated into viral chromatin in replication foci, we performed ChIPseq on growing and differentiated CIN612-9E cells using antibodies against macroH2A1.2. Alignment of the paired-end reads to the HPV31 genome showed binding of macroH2A1.2 across the viral genome, with most binding observed at the 3’ end of the late region and 5’ end of the upstream regulatory region (URR) (Figure 3, upper panel) in both growing and differentiated cells. The relative binding of macroH2A1.2 to the viral genome increased in differentiated conditions relative to the growing conditions. (Figure 3, upper panel). To confirm this, we calculated the binding of macroH2A1.2 to the viral genome using q-PCR. Primers pairs were designed to amplify different regions across HPV31 genome as shown in Supplementary Figure 4A. Chromatin was prepared from growing and differentiated conditions and ChIP was carried out with macroH2A1.2, Histone H3 and IgG antibodies. In support of the ChIP-PCR data, binding of macroH2A1.2 was observed to all regions of the HPV31 genome and with increased peaks under differentiated conditions (Supplementary Figure 4B and 4C). Specifically, upon differentiation macroH2A1.2 binding per viral genome increased 5.2, 5.3, 7.0, 9.3, 10.4, and 3.9 -fold at URR1, URR2, early promoter, early and late region, and L1, respectively (Supplementary Figure 4B and 4C).

**Figure 3.**
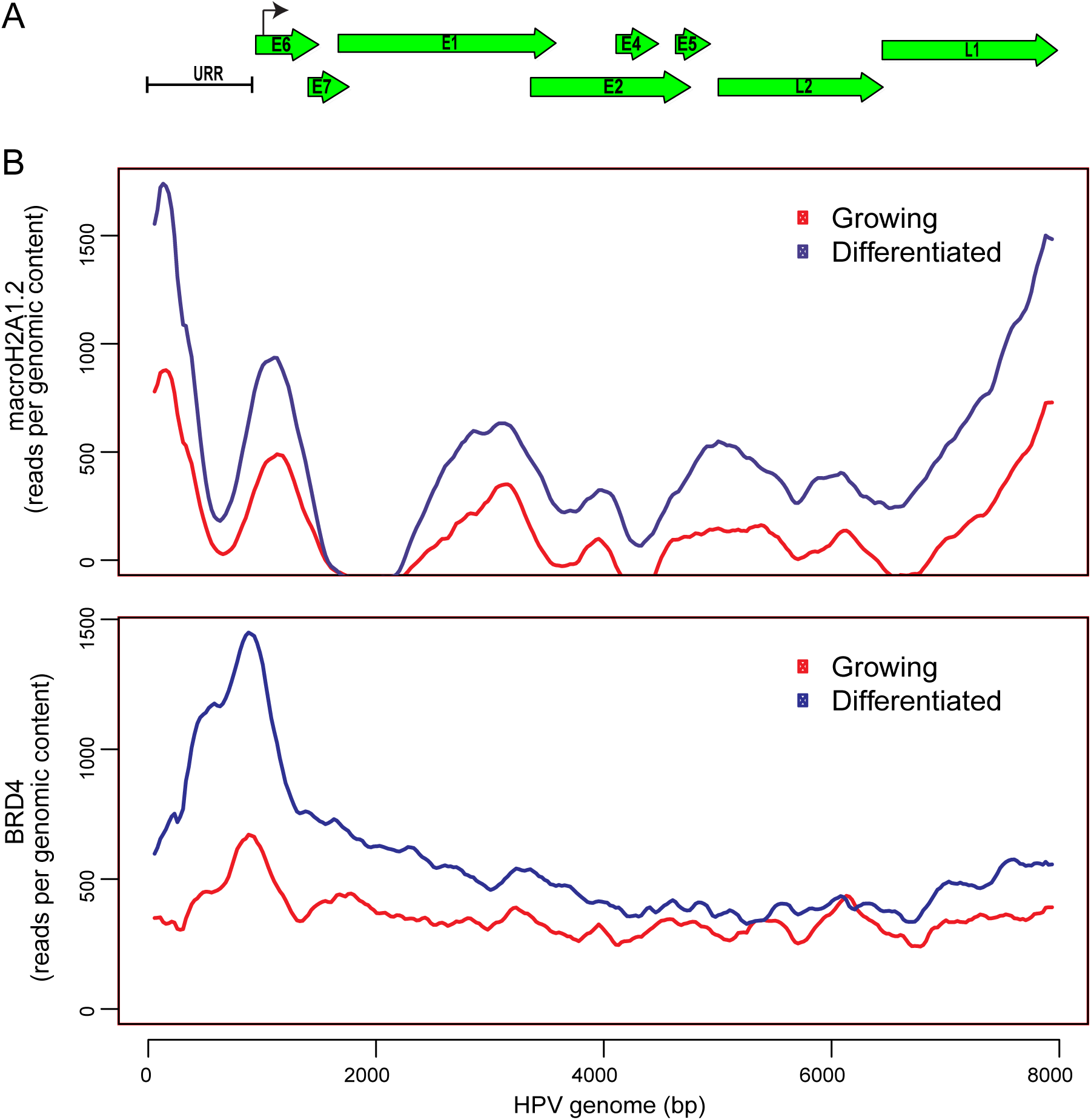
macroH2A1 and Brd4 binding to HPV31 genome is increased in differentiated conditions. **A.** Schematic of a linearized HPV31 genome. URR, upstream regulatory region. **B.** ChIPseq was performed with macroH2A1.2 and Brd4 antibodies. Alignment of ChIPseq reads to the HPV31 reference genome in samples from growing and differentiated 9E cells is shown. ChIPseq reads were aligned and analyzed. Data was averaged from two biological replicates.

We have previously shown that Bromodomain-containing protein 4 (Brd4) is recruited to HPV viral replication factories (29). Brd4 is a chromatin adaptor protein that binds acetylated lysine residues on histone tails and play an important role in transcription (30, 31). Brd4 acts an a scaffold for the assembly of large protein complexes on hyper acetylated promoters and enhancers to promote RNA polymerase II activity to mediate transcription initiation and elongation. For comparison, we carried out ChIPseq to determine the recruitment of Brd4 to the viral genome in growing and differentiated CIN-612 9E cells. Similar to macroH2A1.2 binding, Brd4 recruitment was increased in differentiated cells as compared to cells cultured in proliferative conditions (Figure 3, lower panel). As expected, the transcription modulator Brd4 was enriched primarily at the early enhancer/promoter region of the viral genome.

### macroH2A1 depletion does not affect viral genome amplification in differentiated cells

To analyze the role of macroH2A1 on HPV31 genome amplification, we downregulated both isoforms of macroH2A1 in 9E cells using siRNA and analyzed the viral DNA levels by qPCR and southern blotting. Cells were transfected with Ctrl siRNA or siRNA against macroH2A1 and total cell DNA was isolated at (T=0 days, growing cells) and at (T=8 days, differentiated cells) as shown in the scheme in Figure 4A. The efficiency of macroH2A1 downregulation was determined by Western blotting (Supplementary Figure 2). There was no change in viral copy number in differentiated cells in the macroH2A1 depleted samples as compared to control cells (Figure 4B). Viral genome amplification was also measured by Southern blotting in siCtrl and simacroH2A1 transfected cells. Similar to the data shown in Figure 4B, there was no difference in viral amplification between macroH2A1 depleted and control cells (Figure 4C, left and right panels). There was also no change in the ratio of supercoiled monomeric genomes and higher forms that represent multimeric genomes and/or replication intermediates. Taken together, these data show that the absence of macroH2A1 does not affect HPV31 DNA replication or genome copy number.

**Figure 4.**
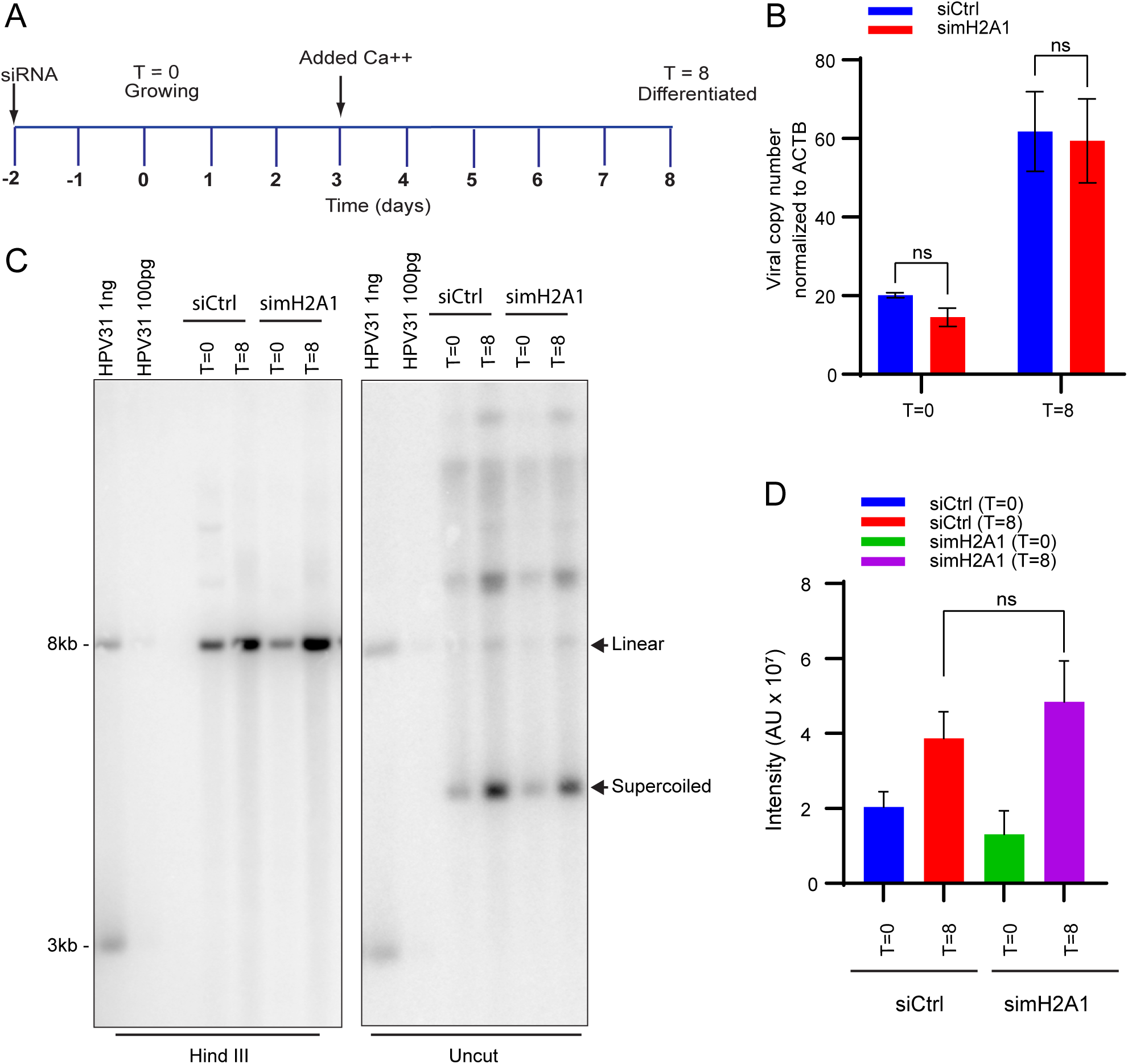
macroH2A1 does not regulate levels of viral replication of HPV31 genomes in CIN612-9E cells. A. Timeline. Cells were transfected with either Ctrl or MacroH2A1 siRNA 24 hours after of plating. Two days later, DNA was extracted at T=0 days for growing or T=8 days for differentiated conditions as indicated in the diagram. Western blot showing the efficiency of downregulation is shown in Supplementary Figure 2 **B.** Viral copy number was measured by qPCR after macroH2A1 siRNA treatment. **C.** Southern blot analysis of DNA extracted from the siCtrl and simacroH2A1 samples after macroH2A1 depletion. The arrows indicate linear and supercoiled viral genomes. The blot is representative of two independent experiments. **D.** Quantification of the linear HPV DNA from the southern blot. Statistical significance was determined using paired t-test for changes in HPV replication. ns: not statistically significant. For B and D, errors bars represent +/- SEM from four and two independent experiments, respectively.

### Depletion of macroH2A1 reduces levels of viral transcripts in differentiated cells

9E cells can be used to study the different stages of the viral life cycle as late viral transcription can be activated by cellular differentiation (32). To determine the role of macroH2A1 in viral transcription, we analyzed early, intermediate, and late HPV31 viral transcripts after depletion of macroH2A1 by siRNA. Ctrl siRNA or siRNA against macroH2A1 were transfected into CIN612-9E cells 24 hours after plating. For the early stages of the viral life cycle, cells were collected under growing conditions, 48 hours after transfection (T=0). For later stages of the viral life cycle, cells were differentiated in calcium containing medium as indicated in the time line in Figure 5A. mRNA expression levels of the macroH2A1 isoforms and viral transcripts including E6*I (early), E1^E4 (intermediate) and L1 spliced (late) were measured by qPCR (Figure 5B and 5C). Both splice variants macroH2A1.1 and macroH2A1.2 were significantly downregulated (∼95%) under growing (T=0) and differentiated conditions (T=8) (Figure 5B). Depletion of macroH2A1 resulted in no significant change in expression levels of early (E6*I) transcripts. However, mRNA expression levels of both intermediate (E1^E4) and late transcripts (L1 spliced) were significantly reduced in differentiated cells upon depletion of macroH2A1 (Figure 5C).

**Figure 5:**
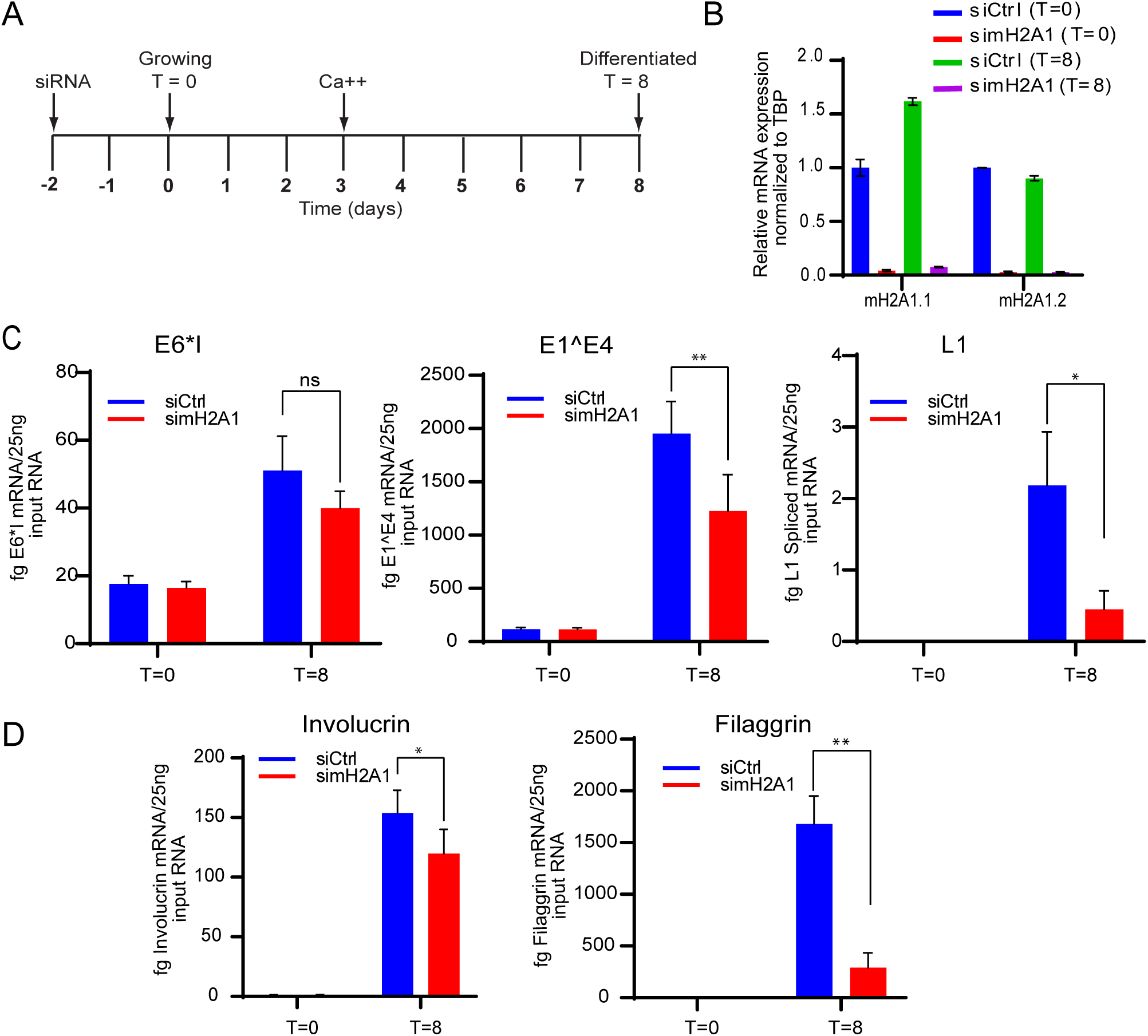
Depletion of macroH2A1 reduces the levels of HPV viral transcripts. **A.** Timeline: 9E cells were transfected with either Ctrl or MacroH2A1 siRNA 24 hours after plating. Two days later, RNA was extracted at T=0 days for growing, or T=8 days for differentiated conditions as indicated in the timeline. **B.** The efficiency of depletion was monitored by measuring levels of macroH2A1.1 and macroH2A1.2 transcripts **C.** Viral transcripts E6*I, E1^E4 and spliced L1 were detected by qRT-PCR **D.** Transcripts for the keratinocyte differentiation markers, involucrin and filaggrin were measured by qPCR, as indicated. All results were obtained from four independent experiments. A paired student t test was used to determine statistical significance. Errors bars represent +/- SEM, *, P<0.05; **P<0.005; ns, not statistically significant.

HPV late gene expression is regulated by the keratinocyte differentiation process and so we monitored this by measuring the mRNA levels of the keratinocyte differentiation induced genes, involucrin and filaggrin (Figure 5D). MacroH2A1 depletion decreased mRNA expression levels of both transcripts in the differentiated cells making it difficult to conclude whether viral transcription was directly regulated by macroH2A1, or whether the reduction in viral transcription was indirect and the result of impaired differentiation. Creppe et al. have shown that macroH2A1 expression increases in the differentiated layers of skin, and that depletion of macroH2A1 in human keratinocytes interferes with activation of differentiation genes and the formation of stem cell holoclones (33). We did not observe any consistent change in the expression of macroH2A1 in our calcium differentiated 9E cells (Supplementary 2B), but downregulation of macroH2A did reduce RNA levels of involucrin and filaggrin. In summary, our data shows that depletion of macroH2A1 reduces HPV31 late gene expression), however this could be indirect due to effects on keratinocyte differentiation.

### MacroH2A1 is not enriched in HPV16 E8^E2 mutant replication foci

Productive replication and transcription of HPV viral genomes is inhibited by the viral E8^E2 repressor protein (16). HPV16 genomes mutated to eliminate expression of E8^E2 have increased viral DNA replication and transcription, even in the absence of keratinocyte differentiation (34). In our laboratory, we find that keratinocyte cell lines containing an HPV16 genome mutated in E8^E2 (HPV16ΔE8^E2) form large replication foci in the absence of keratinocyte differentiation (manuscript in preparation). Therefore, we used cell lines containing wild-type HPV16 and HPV16ΔE8^E2 genomes to analyze the location of macroH2A1 in these replication foci. HPV16 wild-type cells contained only a few, small replication foci. However, in cells containing the HPV16ΔE8^E2 genome, multiple small and large foci were present in the nucleus of many cells (Figure 6A and B). Foci formed with HPV16ΔE8^E2 showed two phenotypes: in ∼77% foci macroH2A1 was mostly depleted; in ∼23% of the foci a small amount of macroH2A1.2 was present as an intensely stained central core (Figure 6A). This showed that macroH2A1.2 was enriched in the small wild type foci but was mostly depleted from the majority of replication foci in HPV16ΔE8^E2 mutant cells (Figure 6B), irrespective of the size. This led us to postulate that the E8^E2 repressor may be involved in the recruitment of macroH2A1 to the replication foci. To ensure that the HPV16ΔE8^E2 replication foci were not simply depleted of all histones, the presence of H2A, H2B, H3 and H4 were analyzed by immunofluorescence in HPV16ΔE8^E2 replication foci. The canonical histones were neither depleted nor enriched in the HPV16ΔE8^E2 replication foci (Supplementary Figure 5).

**Figure 6:**
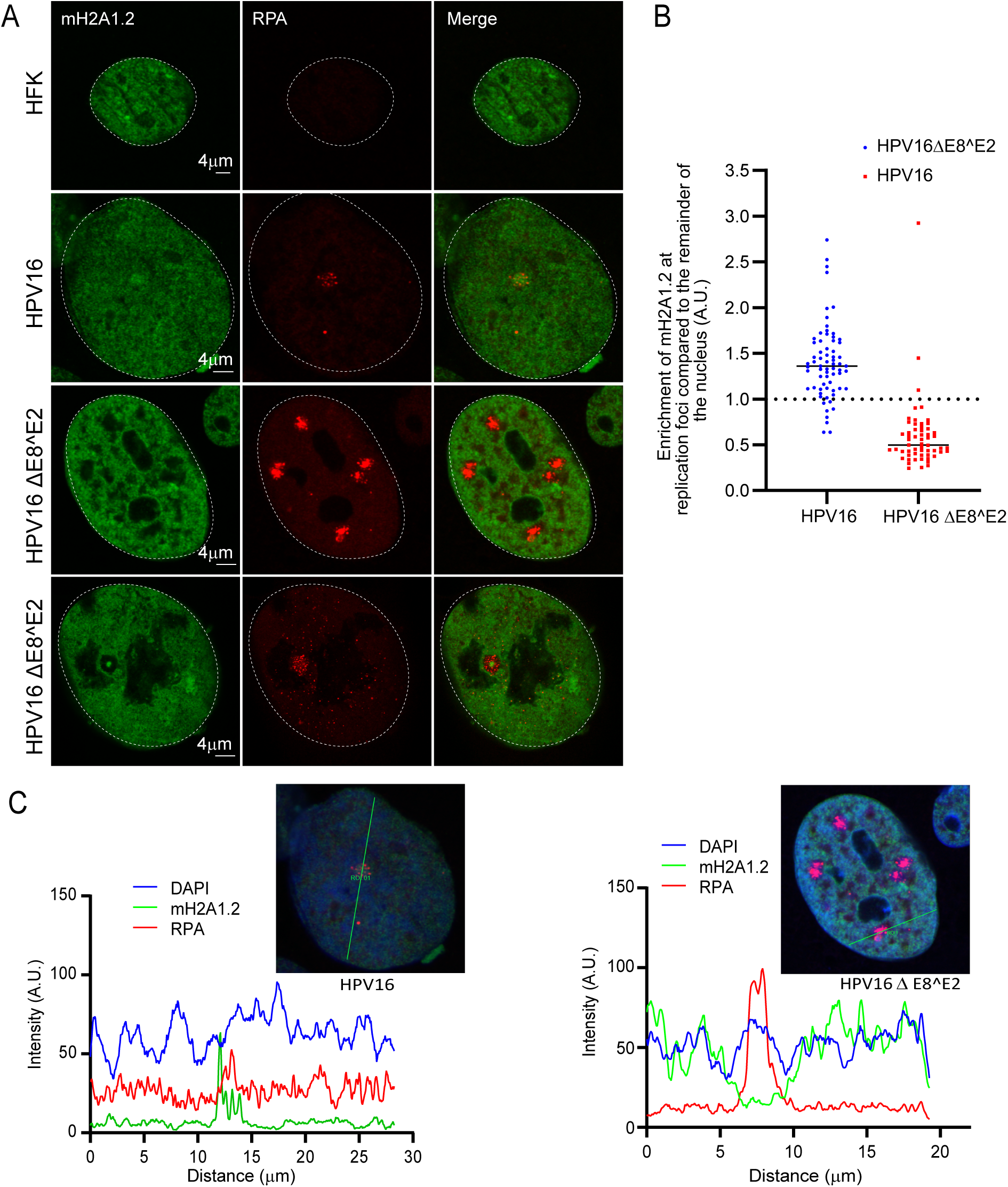
macroH2A1 is not recruited to the replication in HPV16ΔE8^E2 cells. **A.** HFK (strain 20) cell lines containing either HPV16 wild-type or HPV16ΔE8^E2 genome were immunostained with antibodies against macroH2A1.2 (green) and RPA (red). Nuclei are indicated with dotted lines. A white arrow indicates core staining of macroH2A1.2. **B.** Quantification of panel A. The mean fluorescent Intensity of macroH2A1.2 was calculated using ImageJ (values >1.0 indicate enrichment). A total of 136 cells were scored with the parental HFK20 strain as a negative control (no RPA foci were detected). In HPV16ΔE8^E2 cells, 58 RPA positive foci were counted from 41 cells (N=41) in two independent experiments. 94.8% of foci (55/58) showed no enrichment of macroH2A1. In wild-type HPV16 genome containing cells (N=66), 68 RPA positive foci were scored and 89.1% (61/68) showed enrichment of macroH2A1 (mostly single foci are present in wild type HPV16 cells). **C.** Fluorescence intensity light scans obtained by drawing a line through a nucleus shown in panel A.

### MacroH2A1 is incorporated in viral replication foci independent from DNA damage

MacroH2A1.2 is known to mediate a dynamic transition of chromatin from a relaxed accessible state to a condensed inaccessible state at double strand DNA breaks (DSBs) (26). The accumulation of macroH2A1.2 in this compact chromatin environment is ATM-dependent and promotes the retention of the homologous recombination protein BRCA1 at the DSBs. HPV recruits several repair factors from ATM and ATR signaling pathways to replication foci (5, 29, 35) and the enrichment of HR factors indicates that amplification of viral genomes during productive infection might involve recombination-directed replication (5, 11, 12, 36). Therefore, we asked whether macroH2A1 enrichment at viral replication foci correlated with DNA damage signaling. We investigated the localization of mH2A1.1 along with DNA damage markers γH2AX, 53BP1, BRCA1 and RAD51 in the HPV16 wild-type and HPV16ΔE8^E2 mutant cell lines. γH2AX was present in 100% HPV16 wild-type and HPV16ΔE8^E2 replication foci indicating that DNA damage response (DDR) signaling pathways were intact in HPV16 wild-type and HPV16ΔE8^E2 mutant cells (Figure 7A-B). However, despite active DDR signaling in the foci in E8^E2 mutant cell lines, macroH2A1.1 was depleted in ∼94 % of these foci (Figure 7B). BRCA1, RAD51 and 53BP1 were also enriched at 100% of viral replication foci in both HPV16 wild-type and HPV16ΔE8^E2 mutant cells. Altogether, these data show that the enrichment of macroH2A1 at viral replication foci does not correlate with DNA damage signaling.

**Figure 7:**
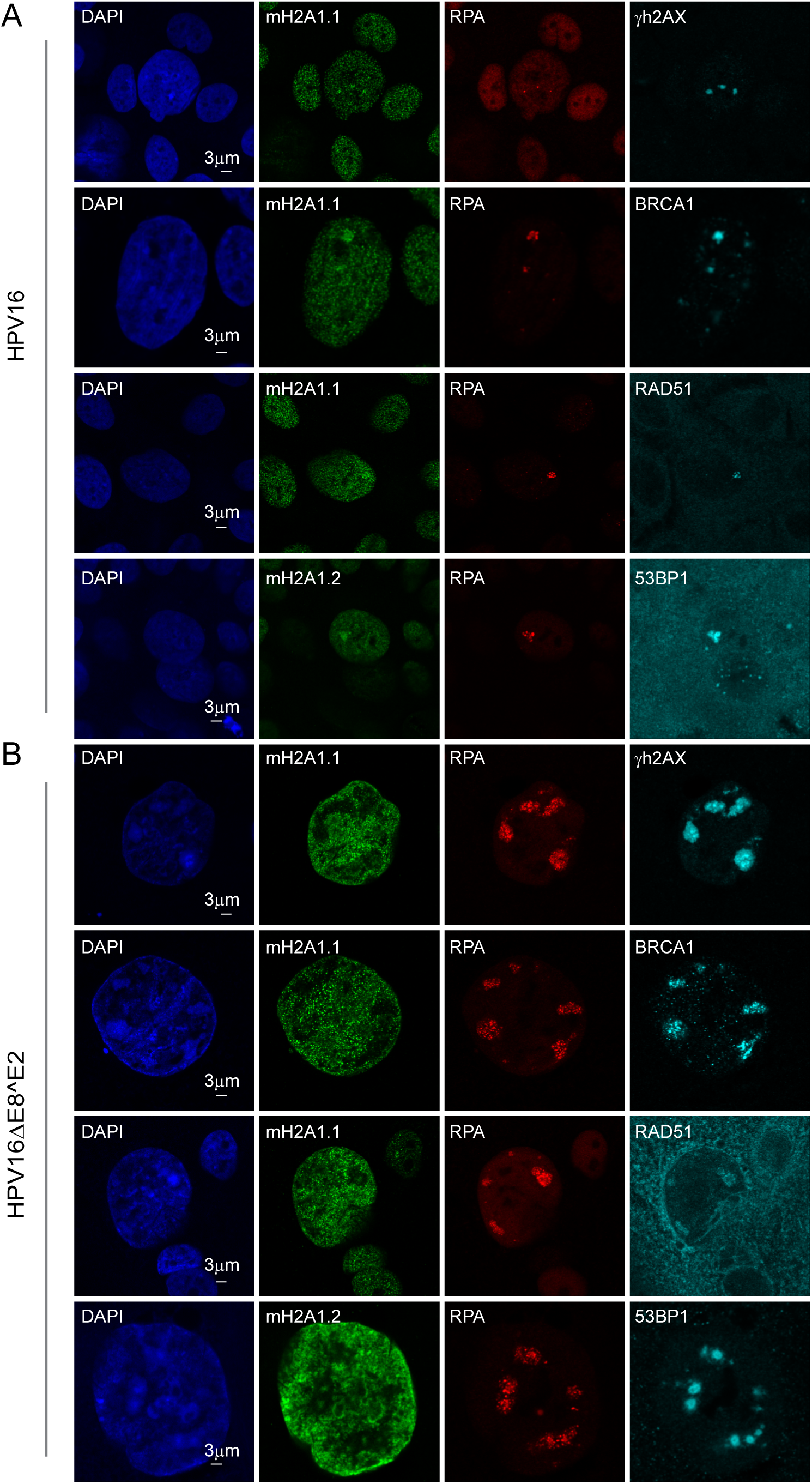
macroH2A1 is recruited to the viral replication foci in a DNA damage independent manner. HFK39 (39 strain) cell lines containing HPV16 wild-type or HPV16ΔE8^E2 genome were immunostained with antibodies against macroH2A1.1 or macroH2A1.2 (green), RPA (red) and several DNA damage associated proteins (cyan). **A.** HFK39 (39 strain) cell lines containing HPV16 wild-type genomes were immunostained with antibodies against macroH2A1.1 or macroH2A1.2 (green), RPA (red) and DNA damage and repair associated proteins (γH2AX, BRCA1, RAD51 and 53BP1 in cyan). For macroH2A1.1 and γH2AX staining, 40 RPA positive foci were scored from 26 cells (N=26) and ∼85 % of foci (34/40) showed enrichment for macroH2A1.1 and 100% for (40/40) γH2AX. For macroH2A1.1 and RAD51 staining in total of 9 cells (N=9), 9/9 RPA positive foci showed enrichment for macroH2A1.1 and (9/9) for RAD51. In total of 20 cells (N=20), 34 RPA positive foci were scored and 100% (34/34) showed enrichment for macroH2A1.2 and 100% (34/34) for 53BP1. Quantitation was from two independent experiments. **B.** HPV16ΔE8^E2 containing cell lines were immunostained with antibodies against macroH2A1.1 or macroH2A1.2 (green), RPA (red) and DNA damage and repair associated proteins (γH2AX, BRCA1, RAD51 and 53BP1, in cyan)). In HPV16ΔE8^E2 cells (N=31), ∼ 93 % of total foci (131/141) showed selective depletion of macroH2A1.1 from the replication foci and 100% (141/141) showed enrichment of γH2AX. In HPV16ΔE8^E2 cells (N=16), 59 RPA positive cells were scored and 91.52 % of foci (54/59) showed depletion of macroH2A1.1 and 100% showed (59/59) enrichment of BRCA1. In HPV16ΔE8^E2 cells (N=18), 80 RPA positive foci were scored from total of 18 cells and 78.3 % (47/60) showed depletion of macroH2A1.1 and 100% (60/60) showed enrichment of RAD51. In HPV16ΔE8^E2 cells (N=33), 115 RPA positive foci were scored and ∼89% (102/115) showed depletion of macroH2A1.2 and 100% (115/115) showed enrichment of 53BP1. All the experiments were quantitated from two independent experiments.

### Cellular RNA transcriptional machinery localizes outside viral replication foci

It has been shown previously that the presence of macroH2A in chromatin inhibits transcription by interfering with transcription factor binding, and SWI/SNF remodeling (37) which correlates with repressive H3K9me3 and H3K27me3 marks. Although, macroH2A1 is predominantly found in repressed chromatin, it can also activate expression of a subset of genes (25). Specifically, macroH2A1 is a positive regulator of a subset of the genes that contain macroH2A1 in the transcribed region. RNA Polymerase II (RNA Pol II) mediated transcription, from initiation to termination, is a highly complex process (38). The C-terminal domain (CTD) of RNA Pol II contains heptad repeats that become phosphorylated at serine position 5 (Ser 5) and serine position 2 (Ser 2) during transcription initiation and elongation, respectively (38). To date, nothing is known about the intranuclear location of HPV transcription with respect to replication foci at late stages of infection. Therefore, we examined the location of RNA Pol II in 9E cells to determine whether it correlated with the enrichment of macroH2A1 at the HPV31 replication foci.

As shown in Figure 8A and 8B, RNA Pol II phosphorylated on serine 2 was observed in two different locations with respect to the replication foci; it was either reduced or absent from the interior of the viral replication foci compared to the rest of the nucleus (∼73% foci) or was present at the periphery in a satellite pattern in a fraction of the foci (∼18% foci). The remainder of the foci (9%) did not show any exclusion of RNA Pol II Ser 2. Figure 8A shows a high-resolution confocal image for RNA Pol II Ser 2 on the surface of the viral replication foci. The Pearson’s coefficient obtained from colocalization analysis confirmed that the localization of macroH2A1.2 and RNA Pol II Ser 2 were mutually exclusive at the replication foci (Figure 8C). Similar to RNA Pol II Ser 2, RNA Pol II Ser5 was detected in a satellite localization in ∼24% replication foci and was reduced or absent inside ∼64 % foci (Supplementary Figure 6). The remaining 12% of foci did not show any exclusion of RNA Pol II Ser5 .

**Figure 8:**
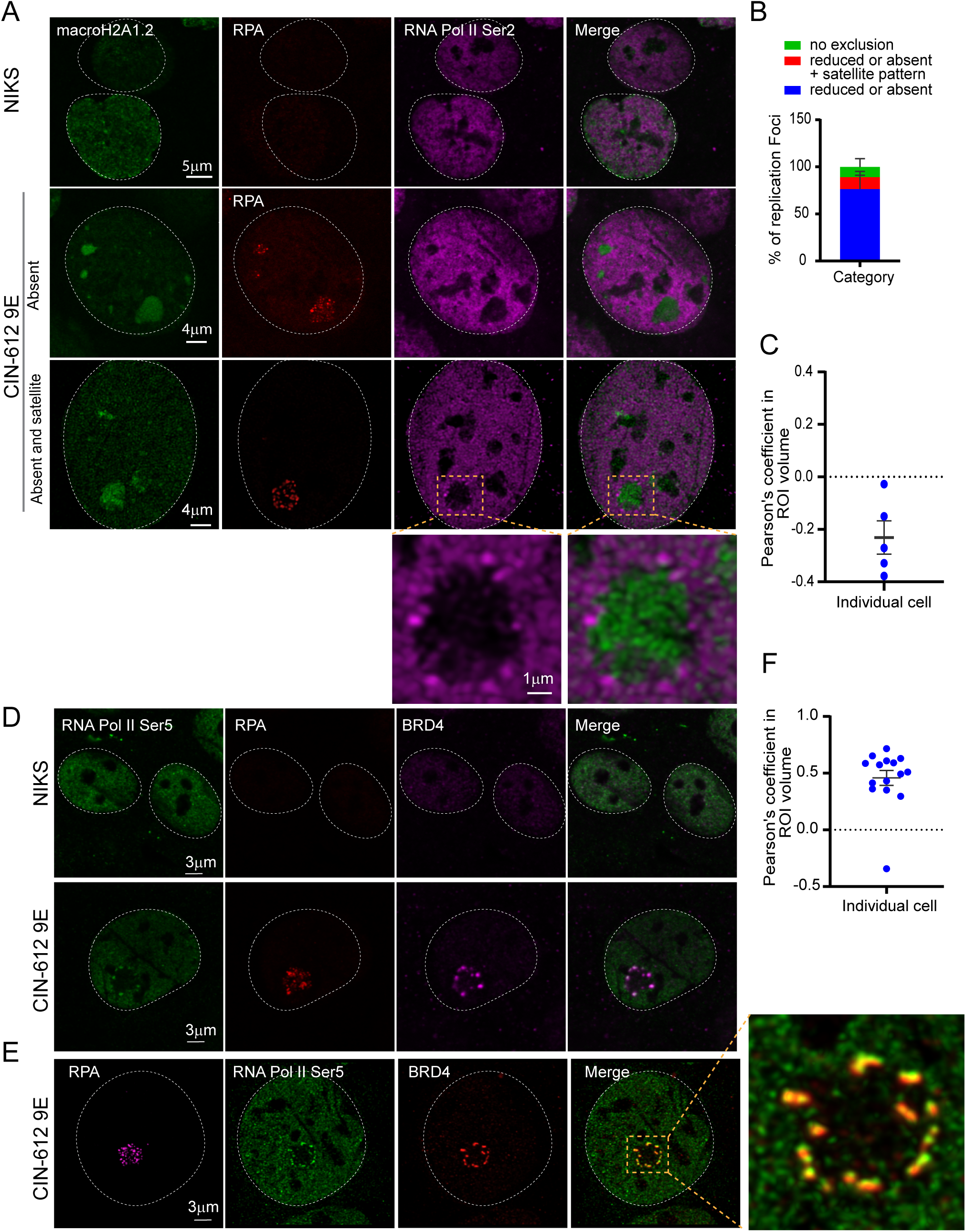
RNA Pol II Ser 2 is located outside the HPV31 replication foci in 9E cells. **A.** Differentiated NIKS or 9E cells were immunostained with antibodies against macroH2A1.2 (green), RPA (red), and RNA Pol II Ser2 (purple). A white dotted line outlines the nuclei. **B.** Distribution of RNA Pol II Ser2 at replication foci by visual counting (N=55 cells, 130 foci) in two independent experiments. **C.** Confocal images (3D) were deconvolved using Huygens Essential. Colocalization analysis was performed in a ROI (region of interest) defined by the RPA signal in Imaris (9.6.0). The % ROI macroH2A1.2 and % of ROI RNA Pol II Ser 2 colocalized were calculated and the Pearson coefficient in the ROI volume was calculated and shown. **D.** Cells were immunostained with antibodies against RNA Pol II Ser 5 (green), RPA (red), Brd4 (CW152 - recognizes both Brd4S and Brd4L; purple). In differentiated CIN-612 9E cells, 233 foci in 71 cells (N=71) were counted using RPA as a marker for viral replication foci. 34 (N=34) cells were scored for differentiated NIKS as a negative control. A white dotted line marks the nuclei. **E.** A high resolution image generated from deconvolved Z stacks collected throughout the nucleus. A single slice is shown, representing the co-localization of Brd4 and RNA Pol II Ser 5 at the periphery of a viral replication foci. The magnified box demonstrates Pol II Ser 5 and Brd4 localization at viral foci. **F.** Confocal images (3D) were deconvolved using Huygens Essential and colocalization analysis was performed in the replication foci ROI defined by the RPA signal using Imaris from total of 15 cells (N=15, foci=41). The % ROI Brd4 and % of ROI Pol II Ser 5 colocalized were calculated and the Pearson coefficients in the ROI volumes are shown.

Notably, the transcription factor Brd4 also localizes in a satellite pattern around the foci in 9E cells (29). Further analysis showed that Brd4 and RNA Pol II Ser 5 colocalized in the satellite pattern on the periphery of viral replication foci and this was confirmed by high resolution confocal imaging (Figure 8D and 8E). The high degree of colocalization was confirmed by colocalization analysis and the Pearson’s coefficient is shown in Figure 8F. These data suggest that very little viral transcription takes place inside the replication foci, and instead viral transcription occurs on the surface of those foci displaying an enriched satellite pattern of RNA polymerase and BRD4. Thus, we propose that there is a spatial separation of replication and transcription at the productive stage of the HPV infectious cycle.

### RNA transcriptional machinery is localized inside HPV16ΔE8^E2 foci

We hypothesized that macroH2A1 might be localized to the replication foci to prevent transcription of replicating DNA, thereby restricting transcription to the surface of the foci. To test this, we assessed the location of RNA Pol II in foci generated by HPV16ΔE8^E2 mutant genomes, which do not contain enriched macroH2A1. As shown in Figure 9, in HPV16 wild type cells, RNA Pol II Ser 5 is often localized outside the foci (with Brd4) and macroH2A1.2 is enriched throughout the foci, as found for HPV31 foci in 9E cells. In contrast, foci formed with HPV16ΔE8^E2 showed two phenotypes: in ∼75% of the foci macroH2A1 was mostly depleted; in ∼25% of the foci a small amount of macroH2A1.2 was present as an intensely stained core (Figure 9A).Furthermore, the location of RNA Pol II Ser5 and Brd4 correlated inversely with these macroH2A1.2 patterns: when macroH2A1.2 was completely depleted from the foci, Brd4 and RNA pol II Ser5 were present inside; however, when macroH2A1.2 was present in the core then Brd4 and RNA Pol II Ser 5 were localized in a ring around the foci (Figure 9A). In conclusion, this data suggests that the function of macroH2A1.2 is to exclude the transcription machinery at the periphery of the foci.

**Figure 9:**
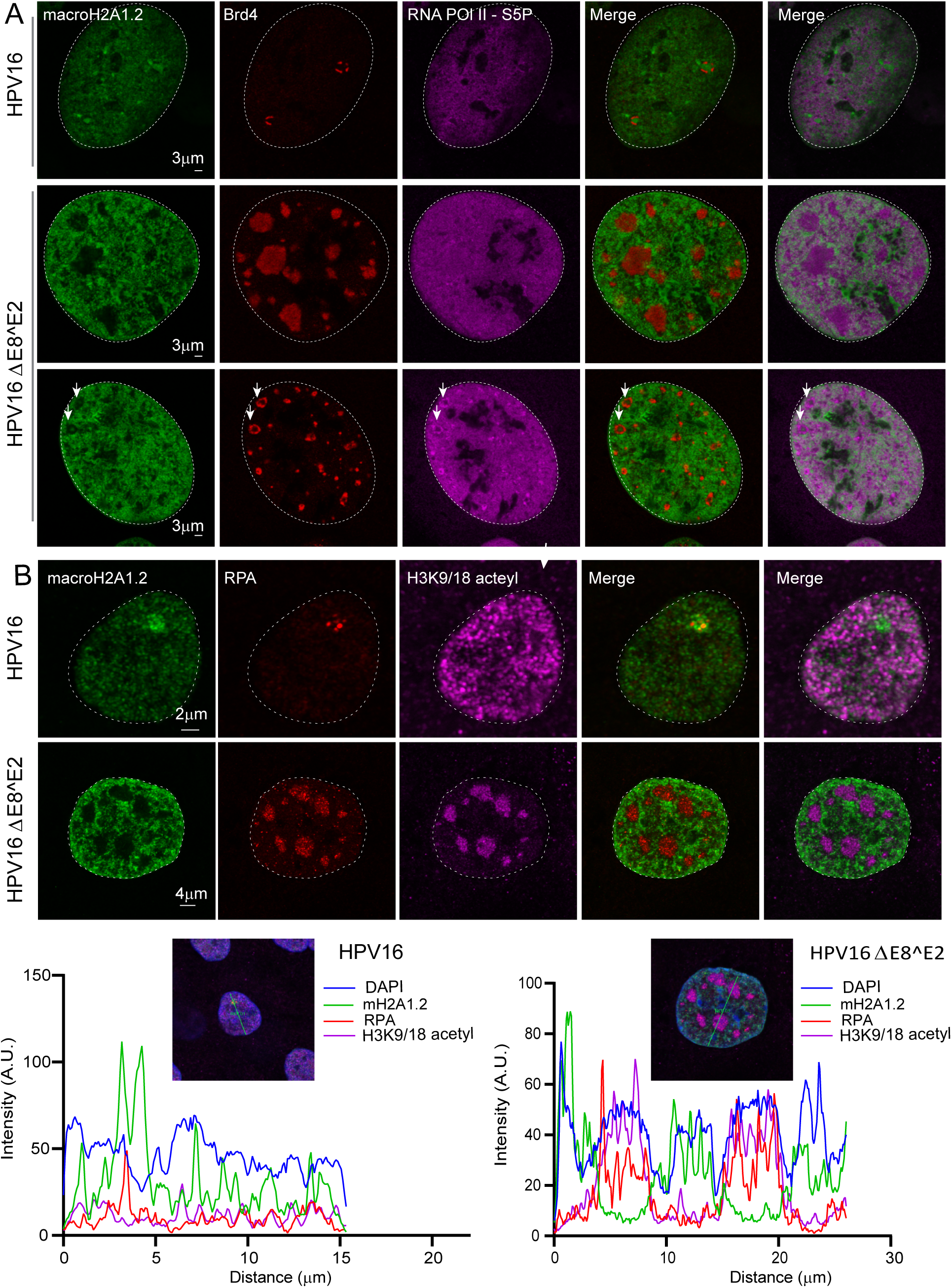
RNA Pol II Ser 5, Brd4 and acetylated histones are localized predominantly inside the replication foci in HPV16 ΔE8^E2 cells. **A**. HFK (39 strain) cell lines containing HPV16 wild-type or HPV16 ΔE8^E2 genomes were immunostained with antibodies against macroH2A1.2 (green), Brd4 (red) and RNA Pol II Ser 5 (purple). Cells containing wild-type HPV16 had RNA Pol II Ser 5 staining outside the foci (∼61% of foci) and in a satellite localization (∼39% of foci). In HPV16 ΔE8^E2 cells, 192 RPA positive foci were scored from total of 32 cells (N=32) in two independent experiments by visual counting. ∼75% of foci (145/192) showed selective depletion of macroH2A1.2 from the replication foci and ∼25% (47/192) showed a residual amount of macroH2A1.2 at the core. Foci indicated with a white arrow show the core staining of macroH2A1.2. RNA Pol II ser 5 and Brd4 were localized inside the foci (∼75% of the foci) and present as a ring in ∼25% of the foci. **B.** HPV16 wild type and HPV16 ΔE8^E2 genome containing cells were immunostained with antibodies against histone H3K9/18 acetyl (purple), macroH2A1.2 (green) and RPA (red). Nuclei were stained with DAPI (indicated by white dotted line) and the distribution of H3K9/18 acetyl in HPV16 wild type (N=35, 37 RPA positive foci) and HPV16ΔE8^E2 cells (N=44, 188 RPA positive foci) were analyzed by visual counting. Cells were scored from two biological independent replicates. Lower Panel: Fluorescence intensity scans obtained by drawing a line through a nucleus using Leica LAS X software.

### MacroH2A1 prevents the formation of acetylated chromatin within replication foci

Brd4 binds to acetylated chromatin through its tandem bromodomains and promotes transcriptional initiation and elongation by recruiting Mediator and pTEFb complexes, respectively (39, 40)). We have also previously shown that the chromatin that surrounds HPV31 foci in 9E cells is enriched in acetylated chromatin (29). To determine the relationship between acetylated chromatin (histone H3 K9ac/18ac), Brd4 and macroH2A1 in the replication foci we analyzed their localization in HPV16 wild-type and HPV16ΔE8^E2 mutant genome containing cells. In support of this hypothesis, macroH2A1.2 is depleted from most of the HPV16ΔE8^E2 mutant foci (∼70% foci) while histone H3K9ac/K18ac is enriched within the foci (Figure 9B). Conversely, macroH2A1.2 was enriched at the HPV16 WT foci, but H3K9/18 acetyl was neither depleted nor enriched at these of the foci (Figure 9B). Altogether, this suggests that the absence of macroH2A1.2 from the foci allows enhanced levels of formation of active chromatin inside the foci.

### The E8^E2 proteins Recruit Co-repressor Complexes to Viral Replication Foci

The E8^E2 proteins from HPV1, 8, 16 and 31 interact with the NCoR/SMRT co-repressor core complex (GPS2, HDAC3, NCoR, SMRT and TBI1 proteins) to repress viral transcription and E1/E2 dependent replication (17). Since macroH2A1 was absent from majority of HPV16ΔE8^E2 foci, we investigated the recruitment of SMRT, along with macroH2A1.2, to HPV31, HPV16 wild-type, and HPV16ΔE8^E2 foci. As shown in Figure 10A (right panel), macroH2A1.2 and SMRT were present in HPV31 and HPV16 wild type foci (∼91 % of foci in 9E cells and ∼68% of foci in HPV16 wild-type cells). In contrast, in the HPV16ΔE8^E2 cell line, macroH2A1.2 was depleted from the foci and there was no enrichment of SMRT. Therefore, there is an association between the enrichment of both SMRT and macroH2A1 to the foci. However, although both factors are present in the foci, they do not localize exactly to the same regions making it unlikely that this is a direct interaction (Figure 10A).

**Figure 10:**
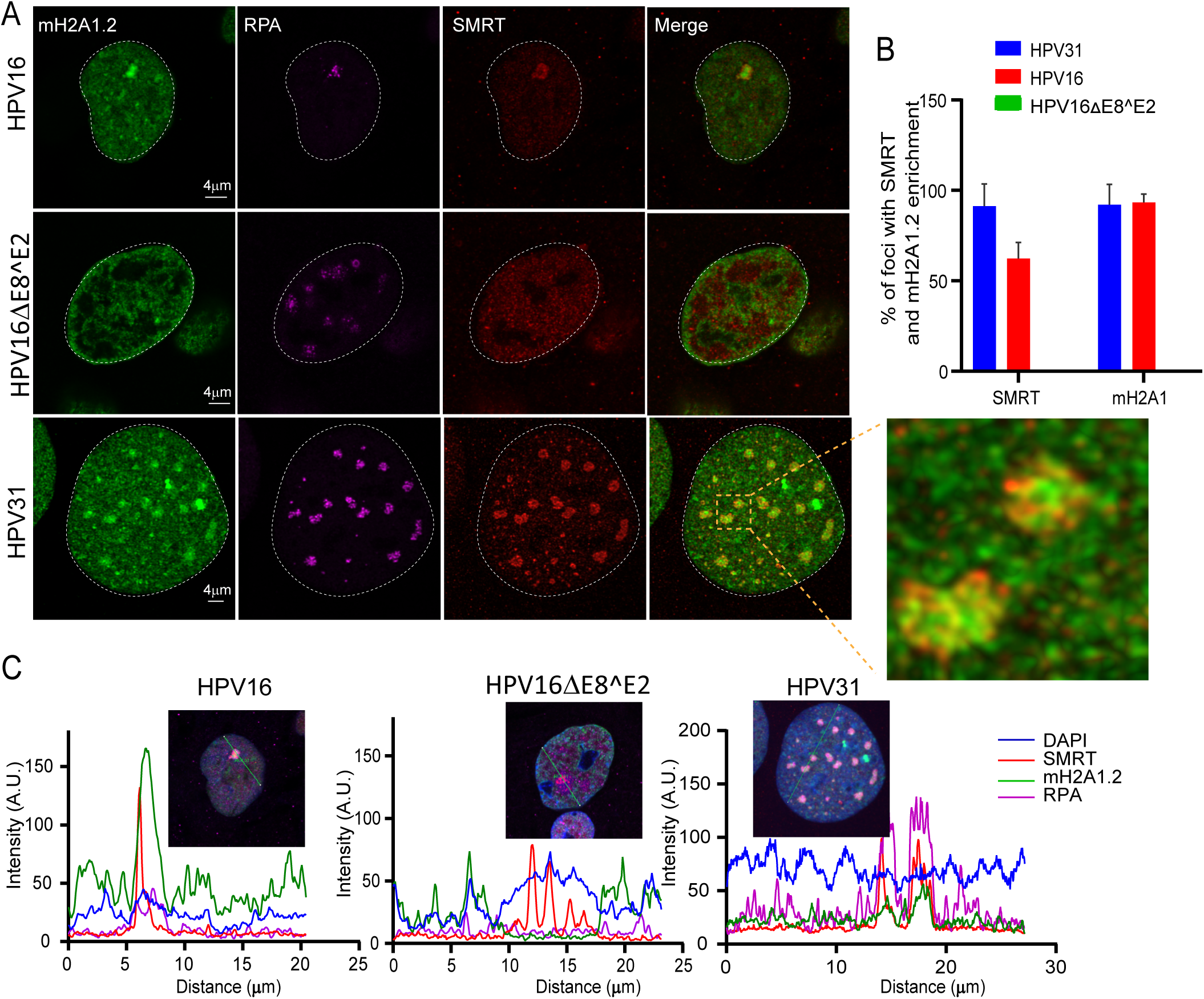
macroH2A1 and SMRT corepressor independently localize to replication foci. **A.** HPV31 (9E cells), HPV16 wild type and HPV16 ΔE8^E2 genome containing cells were immunostained with antibodies against RPA (purple), macroH2A1.2 (green) and SMRT (red). A white dotted line outlines the nucleus. High resolution image of deconvolved image is shown from single slice of Z stacks collected throughout the nucleus for 9E cell. The magnified image in the box area demonstrates macroH2A1.2 and SMRT localization at replication foci. **B.** Percentage of foci enriched for SMRTand macroH2A1.2 in differentiated cells containing HPV31 (N=50 9E cells, RPA positive foci=159), HPV16 wild type (N=28 cells, RPA positive foci=29) and HPV16 ΔE8^E2 (N=53 cells, RPA positive foci=241) were analyzed. Data are from two independent biological replicates. **C.** Fluorescence intensity scans obtained by drawing a line through the nuclei indicated using Leica LAS X software.

### HPV16 E2-TA and E8^E2 proteins are recruited to replication foci in HPV16 wild-type and HPV16ΔE8^E2 genome cell lines

Our observations indicate that transcriptional repressor complexes are present throughout wild-type HPV replication foci and transcriptional activity is restricted to the surface. In contrast, foci generated in the absence of E8^E2 are larger, do not recruit corepressor complexes and contain evidence of transcriptional activity throughout the foci. Dreer et al. have shown that the HPV31 E8^E2 protein localizes to replication foci formed by the E1 and E2 proteins wherein the E8 moiety recruits the corepressor proteins (16). To determine the localization of the HPV16 E2-TA and E8^E2 proteins in the foci we used two different HPV16 E2 antibodies, one that recognizes the unique N-terminal domain of the E2-TA protein and the other that recognizes the DNA binding domain shared by both E2 proteins. As expected, in foci generated by the HPV16ΔE8^E2 genome, the staining pattern was very similar for both antibodies since only the E2-TA protein was present. Two patterns of E2 staining were observed: E2 was either enriched throughout the foci or present in ring form around the foci similar to localization of Brd4 and RNA Pol II Ser5 (Figure 9). This pattern correlated with the macroH2A1 patterns; when macroH2A1.2 was completely depleted from the foci (∼ 85 % of foci), E2 was present inside; however, when macroH2A1.2 was present in the core (∼15% of foci) then E2 was localized as a ring around the foci (Figure 11). In HPV16 wild-type cells, there were three patterns observed. Both antibodies either detected E2 proteins in the satellite regions around the foci (∼22%) or throughout the foci (∼26%). The satellite localization was similar to Brd4 and Pol II Ser 5 localization.However, in most of the cells, only the C-terminal antibody stained throughout the foci (∼52% of the foci); we conclude that this is the E8^E2 protein and its localization is similar to macroH2A1.2 localization (Figure 11).

**Figure 11:**
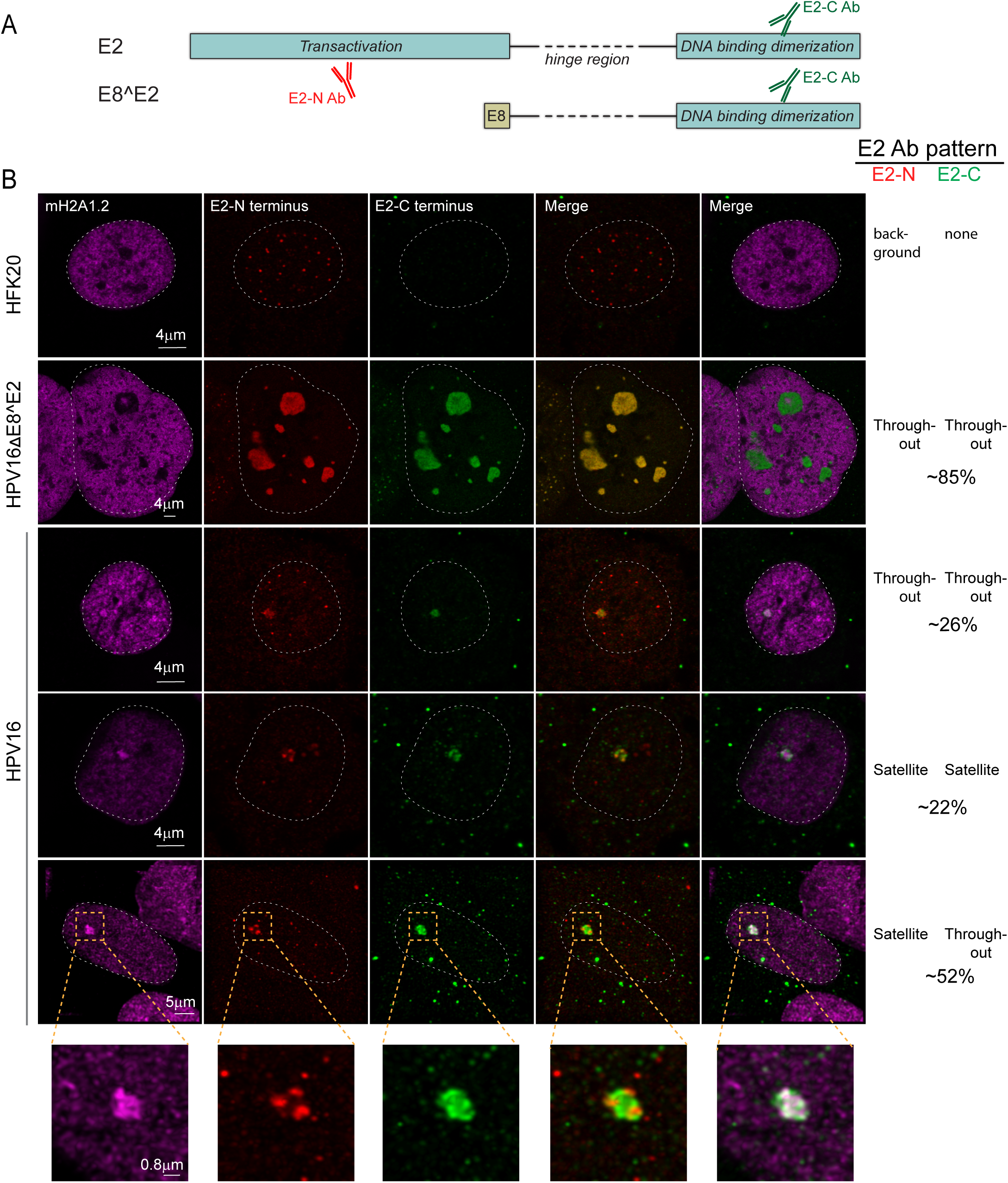
HPV16 E2-TA and E8^E2 proteins are recruited to replication foci in HPV16 wild-type and HPV16ΔE8^E2 genome cell lines. **A.** Diagram of the E2-TA and E8^E2 proteins and antibodies. **B.** HPV16 wild-type and HPV16ΔE8^E2 genome containing cells were immunostained with antibodies against macroH2A1.2 (cyan), E2 (N-terminus antibody; red), E2 (C-terminus antibody; green). A white dotted line outlines the nucleus. In HPV16 wild-type cells, macroH2A1.2 was enriched at ∼100 % (37/37) of foci collected from N=37 cells. Approximately ∼ 52% of foci (19/37) showed a satellite pattern for E2 the N-terminal antibody while the C-terminal E2 antibody stained throughout the foci; ∼ 22% (8/37) off the foci showed satellite pattern for both the E2 N terminus and C-terminus staining pattern; in the remaining ∼26% of the foci (10/37) both antibodies stained throughout the foci. In HPV16ΔE8^E2 genome cell lines, 251 RPA positive foci were scored from N=39 cells and ∼85% of foci (214/251) showed enrichment of E2 (both through N-terminus and C-terminus antibody) inside the foci and (∼15% of the total foci (37/251) showed a ring around the foci for E2 (not shown). Data were obtained by visual counting from two experiments.

Taken together, these studies show that the E8^E2 protein can recruit cellular corepressor proteins to HPV replication foci to repress transcription, while the E2-TA protein is located (at least initially) on the surface of the foci where it colocalizes with Brd4 and RNA Pol II Ser2/5, implying that these are transcriptionally active complexes. The histone variant macroH2A1, is also recruited to foci in cells that express the HPV E8^E2 protein, and we postulate that both cellular macroH2A1 and viral E8^E2 proteins repress transcription of replicating viral DNA. Attempts to detect a direct interaction between these factors have been unsuccessful and the proteins (or their recruited repressors) do not completely colocalize (e.g. Figure 10, macroH2A1 and SMRT). Therefore, we conclude that although macroH2A1 is only recruited to HPV replication foci in cells that express the E8^E2 protein; there is not a direct association.

## Discussion

In this study we show that the variant histone macroH2A is recruited to HPV replication foci and we investigated the potential roles of this protein in the productive stage of the viral life cycle. Using ChIPseq, we find that macroH2A1 is associated with viral DNA and that this association increases in differentiated 9E cells. However, depletion of macroH2A1 in these cells had no effect on HPV31 viral DNA copy number. We also observed that depletion of macroH2A1 reduced the levels of late viral transcripts, but also reduced expression of the keratinocyte differentiation markers (involucrin and filaggrin), indicating that the effect of macroH2A1 depletion on the viral transcription might be indirect. MacroH2A1 is generally bound to repressed chromatin (25, 41); the majority of genes that are oocupied with macroH2A1 are silent. We propose that macroH2A represses transcription of the viral DNA internal to the foci, which is undergoing replication and processing. Our data indicates that viral chromatin on the surface of the foci is transcriptionally active, and so macroH2A could indirectly activate chromatin by organizing the replication foci into transcriptionally active and inactive zones.

For the most part, we observe that, sites of viral replication and transcriptional regulation are spatially separated in HPV infected cells as RPA is localized inside the foci while phosphorylated forms of RNA polymerase, and the transcriptional regulator Brd4 are localized towards the periphery of HPV31 foci. We further propose that the spatial separation of replication and transcription compartments is regulated by association of macroH2A1 with viral chromatin. It has been shown previously that in adenovirus infected cells, replication and transcription sites are spatially separated (42) and that viral replication and transcription are partitioned into different substructures within replication compartments in HSV infected cells (43). Very little is known about the spatial organization of HPV late gene expression and virion assembly and this is a fruitful area for future study.

The presence of Brd4 and the E2-TA protein (in addition to phosphorylated forms of RNA polymerase) on the surface of the replication foci implies that these are regions of transcriptional regulation. Brd4 is most often a positive regulator of transcription, and can activate early HPV transcription (44, 45). However, for the most part Brd4 is a transcriptional repressor in the the presence of the HPV E2 protein (reviewed in (46)). The short form of Brd4 has also been shown recently to repress late viral transcription in undifferentiated cells (47). Nevertheless, the role of Brd4 in regulating viral transcription in productive infection is not well understood and there is evidence that it may activate the late promoter by promoting transcriptional elongation (48). We have attempted to directly detect RNA transcripts within the replication foci using RNA FISH, however the abundance of ssDNA intermediates in the foci has made this inconclusive. Unlike many viruses, HPVs do not induce host transcriptional shut-off and the vast abundance of transcription of cellular genes, in particular rDNA, makes detection of viral transcripts using EU labeling also very difficult. Brd4 has also been shown to ensure efficient transcriptional elongation by preventing R-Loop formation and transcription-replication conflicts (49); this could be very important for a virus that must synthesize large amounts of viral DNA and late mRNA to produce progeny virus.

HPV activates the ATM DNA damage pathway, and this required for viral genome amplification in differentiated cells (5). Various components of the DNA damage response pathway localize to HPV replication factories in differentiating CIN612-9E cells (5, 9, 12, 35). In this study, we have shown that both splice variants of histone macroH2A1, macroH2A1.1 and macroH2A1.2, are recruited to foci, which are sites of HPV DNA synthesis. Moreover, we show that macroH2A1.2 is associated with viral chromatin, and this association is increased during the productive phase of the viral life cycle . MacroH2A1.2 is recruited to sites of double strand DNA breaks and this links the compaction of DSB-proximal chromatin to the BRCA1 dependent homologous recombination repair pathway (26). The dependence of macroH2A1 accumulation on the ATM and ATR signaling pathways at double strand break sites and fragile sites upon replication stress suggested that the DNA damage response pathway could be involved in the recruitment of macroH2A1 to the replication viral foci (50). However, here we show that macroH2A1 is not recruited to the replication foci in a DNA damage dependent manner since the DNA damage associated proteins BRCA1, 53BP1 and RAD51 localize to E8^E2 mutant viral replication foci in the absence of macroH2A1 colocalization.

The E8^E2 protein is a repressor of HPV transcription and replication and it has been shown to localize to E1-E2 dependent replication foci (16, 17). In support of this, we find that both E2-TA and E8^E2 also localize to replication foci generated in differentiated keratinocytes containing HPV16 genomes. Using antibodies against the E2 N-terminal domain (E2-TA only) and C-terminal domain (both E2-TA and E8^E2), we find that the E2-TA protein localizes in a ring around both wildtype and E8^E2 mutant foci, but is later distributed throughout, similar to what we have shown previously in transient replication foci (29). In some wild-type cells, however, we observe E2 within the foci with the C-terminal antibody but not with the N-terminal antibody, indicating that this is the E8^E2 protein. Moreover, the E8-associated corepressor protein, SMRT, also localizes within these foci and we propose that, E8^E2 localizes within the foci to restrict viral transcription. macroH2A1 is also found within the foci in wild-type HPV containing cells but is absent in E8^E2 mutant cells. We have been unable to detect an interaction between macroH2A1 and the E8^E2 protein, and although they are both present in the interior of the replication foci, they do not completely colocalize. Future studies will investigate the relationship between these repressive proteins.

## Materials and Methods

### Cell Lines

CIN612-9E cell line harboring extrachromosomal HPV31 genomes derived from CIN1 HPV31-positive patient biopsy have been described (51). NIKS cells have been described previously (52). Human keratinocytes were isolated from neonatal foreskins as described previously (53). Cell lines containing replicating viral genomes were generated using HPV18 minicircle genomes (54), or either wildtype or E8^E2 mutant HPV16 genomes (55). HPV genomes were generated either by minicircle technology (54), or by removing the vector by restriction digestion and religation, and electroporated into NIKS or HFKs with a pRSV2neo plasmid. Cells were selected for five days with 200µg/ml G418, and cultured until colonies formed, at which point cells were pooled. Cells were checked by Southern blot analysis for extrachromosomal viral genomes before use.

### Cell Culture

All cells were cultured in Rheinwald Green F medium (3:1 Ham’s F12/DMEM-high glucose, 5% fetal bovine serum, 0.4 µg/ml hydrocortisone, 8.4 ng/ml cholera toxin, 10 ng/ml epidermal growth factor (EGF), 24 µg/ml adenine and 6 µg/ml insulin) on lethally irradiated NIH J2 3T3 murine fibroblasts. NIH J2 3T3 mouse fibroblast cells were cultured in DMEM containing 10% newborn calf serum. Mouse feeders were exposed to 60 grays of γ-irradiation before co-culture with keratinocytes. For differentiation, keratinocyte cell lines were cultured in F medium until confluent, and then changed to low calcium basal keratinocyte medium supplemented with SingleQuots (bovine pituitary extract, hydrocortisone, and epidermal growth factor) (Lonza Corporation). After 24 hours, medium was changed to basal medium supplemented with 1.5 mM calcium chloride and cells cultured for five days.

### SiRNA transfections

For transient transfection with siRNA, CIN612-9E cells were plated on lethally irradiated J2/3T3 mouse fibroblasts and cultured overnight. After 24 hours, cells were transfected either with siRNA against non-targeting control (D-00181a0-10-20, Dharmacon) or against macroH2A1 (E-011964-00-0005, Dharmacon). Transfections were carried out according to the manufacturer’s protocol. Briefly, siRNAs were mixed with Lipofectamine RNAiMAX transfection reagent (Thermo Fisher Scientific) in OptiMEM medium and added to cells at a final concentration of 20 nM. Cells were incubated with siRNA for the indicated time periods before removing feeders and harvesting cells for further experimentation.

### Viral genome copy number

Total cellular DNA was isolated from growing or differentiated CIN612-9E cells using the DNeasy Blood and Tissue Kit (Qiagen) at times indicated in the figures. One to five ng of DNA was analyzed by qPCR using 300 nM primers and SYBRgreen Master mix (Roche AG) using a QuantStudio 7 Flex Real Time PCR System (Applied Biosystems). Primers are listed in Supplementary Table 1.

### Southern blot analysis

Total DNA was isolated from CIN612-9E cells using the DNeasy Blood and Tissue Kit (Qiagen). DNA (2μg) was digested with either a single-cut linearizing enzyme (HindIII) for the HPV genome, or with a non-cutter (BamHI) to linearize cellular DNA. After digestion, samples were separated on an 0.8% agarose-TAE gel and transferred onto nylon membranes using a Turbo Blotter (GE Healthcare). Membranes were UV cross-linked (120 mJ/cm^2^), dried and prehybridized in hybridization buffer (3X SSC, 2% SDS, 5X Denhardt’s solution, 0.2 mg/ml sonicated salmon sperm DNA) for one hour. The membrane was hybridized overnight with 25ng (^32^P)-dCTP labeled HPV31 DNA probe in hybridization buffer. The membrane was washed in 0.1% SDS/0.1X SSC and hybridized DNA was visualized and quantitated by phosphor-imaging on a Typhoon Scanner (GE Healthcare). The ^32^P-radiolabeled probe was generated from a plasmid containing the entire HPV16 genome by radiolabeling using a Random Prime labeling kit (Roche).

### RNA extraction and qPCR of viral and cellular transcripts

Total RNA was extracted using RNeasy Mini-RNA extraction kit (Qiagen). RNA concentration was determined using a Nanodrop 1000 spectrophotometer (Life technologies). RNA integrity was checked by capillary electrophoresis on a 2100 bioanalyzer system using RNA 6000 nano kits (Agilent Technologies). Reverse transcription reactions were carried out with Transcription First-Strand Synthesis kit (Roche AG) according to the manufacturer’s protocol. mRNA expression of the indicated genes was analyzed by QuantStudio 7 Flex Real Time PCR System (Applied Biosystems) using SYBR green master mix. Cloned cDNA plasmids (2.5X10^5^-2.5X10^-2^ fg) were included in each run to generate a standard curve. Cloned cDNA standards for the HPV31 spliced mRNAs including E1^E4 (nt 857-877^3292-3296), E*I (nt 186-210^413-416) and L1 (nt 3562-3590^5552-5554) and for the cellular genes involucrin, filaggrin and cyclophilin A were described previously (56). The relative expression of macroH2A1.1 and macroH2A1.2 was calculated by 2−(ΔΔCt) method. The primers used are listed in Supplementary Table 1, and were described previously (26).

### Indirect Immunofluorescence (IF) and image processing

CIN612-9E or HFKs were cultured on glass coverslips and fixed with 4% paraformaldehyde in PBS. After fixation the cells were permeabilized with 0.5% Triton X-100 in PBS and blocked in 5% (v/v) normal donkey serum (Jackson Immunoresearch). Cells were incubated with primary antibodies at 37°C for 1 hour. Primary antibodies used for IF were : macroH2A1.1 rabbit monoclonal (Cell signaling, 12455, dilution 1;100), macroH2A1.2 mouse monoclonal (Millipore, MABE61, dilution 1:100), H2A rabbit polyclonal (Abcam, ab 18255, dilution 1:100), H3 rabbit polyclonal (Abcam, ab 1791, dilution 1:100), H4 rabbit polyclonal (Abcam, ab10158, dilution 1:100), RPA rat monoclonal (Cell Signaling, 2208, dilution 1:250), γH2AX mouse monoclonal (Millipore, 05-636, dilution 1:100), BRCA1 mouse monoclonal (Santa Cruz sc-6954, dilution 1:100), RNA Pol II Ser -2 rabbit monoclonal (Abcam, ab 5095, dilution 1:100), RNA Pol II Ser 5 mouse monoclonal (Abcam, ab5408, dilution 1:100), H3 acetyl K9/18 rabbit polyclonal (Upstate (Millipore), dilution 1:100), E2 HPV16 N-terminus sheep polyclonal (Iain Morgan/Joanna Parish, dilution 1:200), SMRT rabbit polyclonal (Bethyl laboratories, dilution 1:100), Brd4 (CW152, dilution 1: 300), (29)). The macroH2A1.1 and macroH2A1.2 antibodies were validated by depleting macroH2A1.2 using siRNA transfection (Supplementary Figure 2). After primary antibody incubation, the cells were washed three times with PBS and incubated with the secondary antibodies (Alexa 488, Alexa 594, Rhodamine Red-X and Alexa 647 conjugated to the target species (Jackson Immunoresearch) were added at 37°C for 40 minutes. Nuclei were stained with DAPI and coverslips were mounted using 10 μl of prolong Gold (Life Technologies). All Images were acquired with Leica TCS-SP5 or TCS-SP8 laser scanning confocal microscopes (Leica Microsystems) using 63X oil immersion objective (NA 1.4). All 2D images were grabbed as a single optical slice for all the experiments unless otherwise mentioned. 3D images were deconvolved using Huygens Essential (version 20.04, Scientific Volume Imaging B.V.) using theoretical PSF and manual background sustraction. All images were processed using LASAF Lite (Leica microsystems) or Imaris (version 9.6.0, Bitplane).

### Combined immunofluorescence and in situ hybridization (IF-FISH)

Paraformaldehyde fixed cells were stained with macroH2A1.2 antibody as described above. After immunostaining, cells were fixed for the second time with methanol and acetic acid solution (3:1 v/v) at room temperature for 10 min followed by fixation with 2% PFA for 1 minute. Coverslips were treated with RNace-iT cocktail (1:1000 dilution, Agilent Technologies) at 37°C for 1 hour and dehydrated in a 70%, 90%, and 100% ethanol series for three minutes each and air dried. DNA FISH probes were prepared using the FISH-Tag DNA Multicolor Kit labeling kit (Life Technologies) according to manufacturer’s protocol. Hybridization was performed overnight in 1X Hybridization Buffer (Empire Genomics) with 50-75ng of labeled probe DNA at 37°C. Slides were washed at room temperature with 1X phosphate-buffered detergent (PBD, MP Biosciences), followed by washing with wash buffer (0.5X SSC, 0.1% SDS) at 65°C. Nuclei were stained with DAPI and coverslips were mounted using Prolong Gold (Life Technologies) and images were captured using Leica TCS-SP5 or TCS-SP8 microscopes (Leica Microsystems).

### Image Analysis

To measure enrichment of macroH2A1 at the viral replication foci, ImageJ (java. version;1.8.0_112) was used to define region of interests (ROIs) around the viral replication foci (around RPA signal) and equivalent non-replication foci regions in the nucleus and cytoplasm. The mean fluorescence intensity within the ROIs was measured for macroH2A1. Background intensity values (cytoplasm) were subtracted, and the ratio of intensity within the replication foci to non-replication foci was calculated. 3D images were collected as noted in the figure legends. 3D images were deconvolved using Huygens Essential (20.04 Scientific Volume Imaging B.V.). All images were processed using LASAF Lite (Leica microsystems) and Imaris (version 9.6.0, Bitplane). For the colocalization analysis, 3D images were first deconvolved in Huygens Essential. Replication foci ROI were defined by RPA signal using surface masking technique in Imaris. Then colocalization analysis and Pearson’s coefficient was then calculated in replication foci using Imaris

### Chromatin Immunoprecipitation

Chromatin from growing and differentiated CIN612-9E cells were prepared as described previously (56, 57). Briefly, following formaldehyde fixation and glycine quenching, cells were resuspended in cell lysis buffer I to isolate nuclei (50 mM Hepes KOH pH 7.5, 140 mM NaCl, 1 mM EDTA, 10% Glycerol, 0.5% NP-40, 0.5% Triton X-100). Lysates were prepared from isolated nuclei as described in Stepp et al., 2017 (56)). Lysates were sonicated using an ultrasonicator water bath (Bioruptor, Diagenode). A Southern blot was performed to ensure the optimal shearing of viral chromatin (200-500bp fragments). DNA shearing was also measured by agarose gel electrophoresis. For immunoprecipitations, 20 μg of chromatin was incubated with either 3 μg of rabbit IgG (ChromPure), macroH2A1.2 (Millipore, MABE61) or BRD4 (Bethyl, A301-985A100) antibodies overnight at 4°C. Immune complexes were collected using 30 µl Protein G Dynabeads (Invitrogen). DNA was purified using ChIP DNA Clean and Concentrator kit (Zymo Research). Immunoprecipitated HPV chromatin was quantified by comparison with an HPV31 plasmid standard curve using QuantStudio 7 Flex Real Time PCR system.

### Chromatin Immunoprecipitation and Illumina sequencing

Chromatin from growing and differentiated CIN612-9E was processed as described above. ChIP and input libraries were sequenced (2 x 150 bp paired end reads) using the illumina HiSeq 4000 platform (Genomics Resource Center, Institute for Genome Sciences, University of Maryland). Reads were trimmed with Cutadapt version 1.18 (58). All reads aligning to the Encode hg19 v1 blacklist regions (59) were identified by alignment with BWA version 0.7.17 (60) and removed with Picard SamToFastq (The Picard toolkit. https://broadinstitute.github.io/picard/). The remaining reads were aligned to an hg19 reference genome with an additional HPV31 chromosome (GenBank ID: J04353.1) using BWA. Reads with a mapQ score less than 6 were removed with SAMtools version 1.6 (61) and PCR duplicates were removed with Picard MarkDuplicates. Replicate ChIP samples were merged after deduplication using SAMtools. Data was converted into bigwigs for viewing and normalized by reads per genomic content (RPGC) using deepTools version 3.0.1 (62) using the following parameters: --binSize 25 --smoothLength 75 --effectiveGenomeSize 2700000000 --centerReads --normalizeUsing RPGC. RPGC-normalized input values were subtracted from RPGC-normalized ChIP values of matching cell type genome-wide using DeepTools with --binSize 25. Here, the human genome reads were used only to normalize the viral read counts and the full analysis of macroH2A binding to the human genome will be published elsewhere.

### Western blot

Media was removed and J2 fibroblast feeder cells were removed with Versene (Thermo Fisher). Keratinocyte monolayers were rinsed with ice-cold PBS. Growing and differentiated cells were lysed on the plate with 1 ml SDS Lysis Buffer (1% w/v SDS, 10 mM Tris-HCl pH 8, 1 mM EDTA pH 8) heated to 95°C. After scraping the plate, samples were transferred to a low protein binding microfuge tubes and sonicated using a Bioruptor (30 seconds on, 30 seconds off, for 6 cycles, at high power). After sonication, samples were heated at 95°C for 10 minutes in a heat block and centrifuged at 16,100 x g for 5 minutes to remove any debris. 10–20 μg total protein was supplemented with 50 mM DTT and 4X LDS sample buffer (Life Technologies). Samples were heated to 70°C for 10 minutes, cooled to room temperature, and proteins separated by SDS-PAGE on 4-12% NuPage gradient gels (Life Technologies). Proteins were transferred overnight onto PVDF membrane. Western blots were performed using the following antibodies; anti-macroH2A1.1 (Cell signaling, 12455, dilution (1:1000) anti-macroH2A1.2 (Millipore, MABE61, dilution (1:1000)), anti-Histone H3 (Upstate Millipore 07-690, dilution (1:10000). Horseradish peroxidase conjugated secondary antibodies (anti-rabbit, Invitrogen 31460, and anti-mouse Invitrogen, 31430) were used at 1:10,000 dilution. The antibodies were detected using chemiluminescent reagents (SuperSignal Dura Western Detection) and the signal detected and quantitated using a G: Box (Syngene).

## ETHICS STATEMENT

Primary human keratinocytes were isolated from anonymized neonatal foreskins provided to the Dermatology Branch at NIH from local hospitals. The NIH Institutional Review Board (IRB) approved this process and issued an NIH Institutional Review Board waiver.

## ACKNOWLEDGEMENTS

We thank all members of the McBride laboratory for helpful discussions and Tom Kristie for comments on the manuscript. This work was supported by the Intramural Research Division of the National Institute of Allergy and Infectious Diseases at the National Institutes of Health.

## SUPPLEMENTARY FIGURE LEGENDS

**Supplementary Figure 1.**
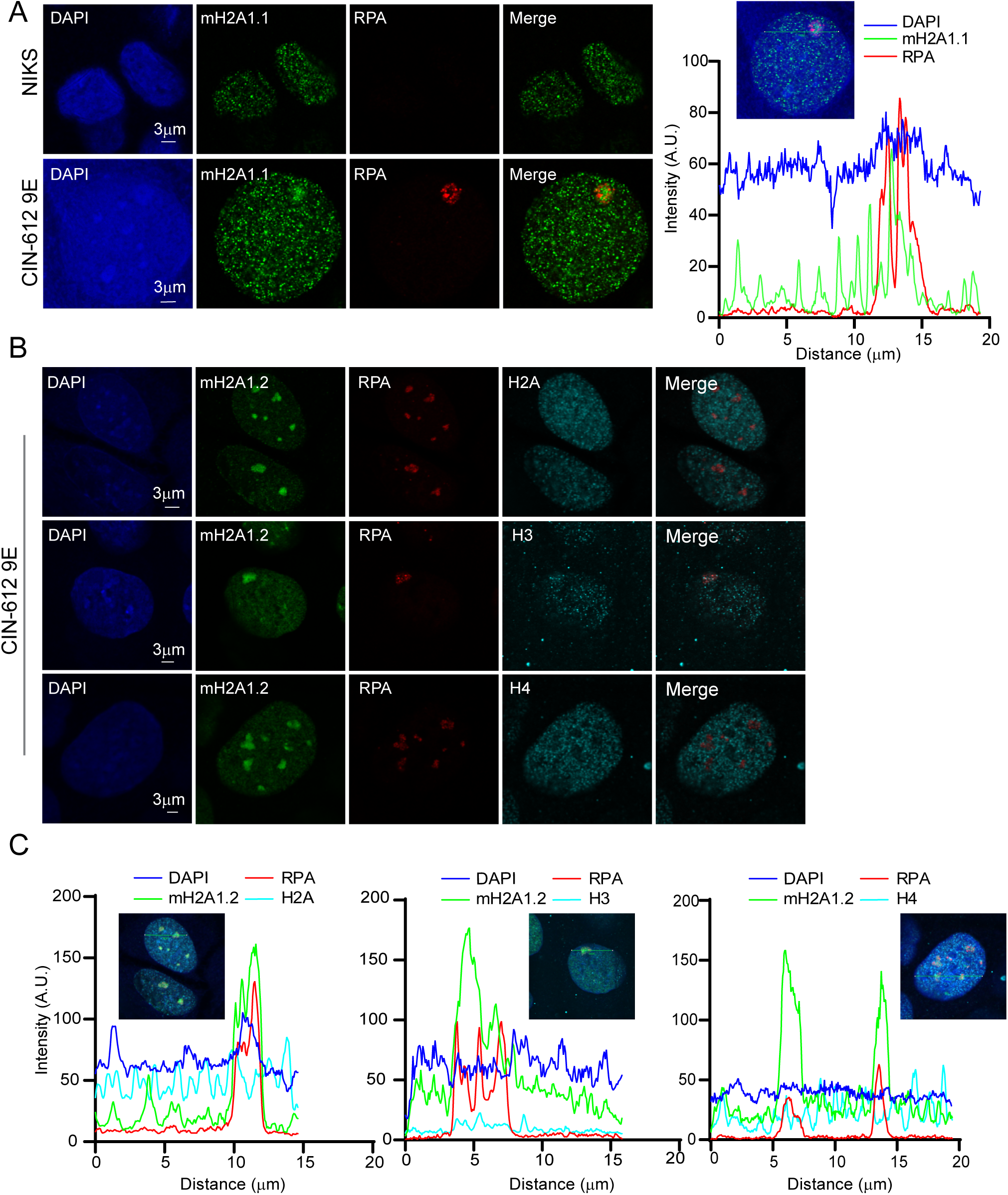
Distribution of macroH2A1.1 and Histone H2A, H3 and H4 in differentiated NIKS and 9E cells. **A.** Cells were immunostained with antibodies against macroH2A1.1 (green), RPA (red) and nuclei were stained with DAPI (blue). In 9E cells (N=47), 129 RPA positive foci were visually scored in two independent experiments and 51% (67/129) showed enrichment for macroH2A1.1. The right Panel shows the fluorescence intensity line scan obtained by drawing a line through the nucleus shown in B using Leica software. **B.** Histone H2A, H3 and H4 are not enriched at RPA foci. Distribution of histone H2A (N=67 cells, N=428 RPA positive foci), histone H3 (N=47 cells, N= 103 RPA positive foci) histone H4 (N=52 cells, N=140 positive RPA foci).Cells and foci were counted from two independent experiments. **C.** Fluorescence intensity line scan obtained by drawing a line through the nuclei shown in B using Leica software.

**Supplementary Figure 2.**
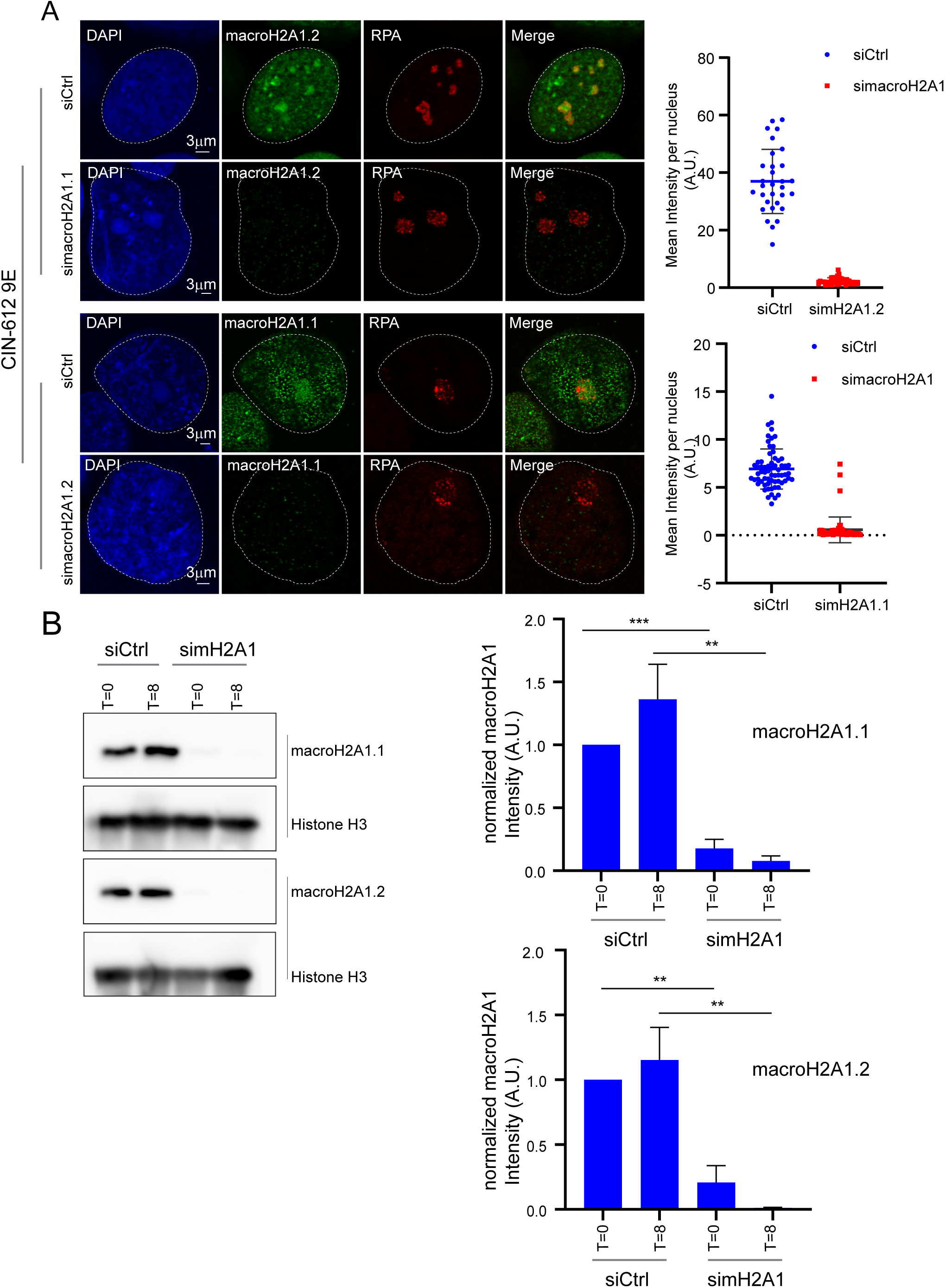
Validation of macroH2A1.1 and macroH2A1.2 antibody in 9E cells. 9E cells were transfected with either Ctrl or macroH2A1 siRNA (targeting both isoforms) 24 hours after plating. Cells were fixed under growing or differentiated conditions as shown in the timeline in Figure 4A. (upper panel) Immunostaining was performed using anti-macroH2A1.2 (green) and RPA (red) antibodies and cells were evaluated for macroH2A1.2 staining in RPA foci (a total of N=31, N=36 in siCtrl and simacroH2A1 cells, respectively) in two independent experiments. (Lower Panel) Immunostaining was performed with anti-macroH2A1.1 (green) and RPA (red) antibodies. Cells were evaluated for macroH2A1.1 staining in RPA foci (N=65, N=55 for siCtrl and simacroH2A1, respectively) in two independent experiments. Nuclei were stained with DAPI (blue). Mean fluorescence Intensity for macroH2A1.1 or macroH2A1.2 per nucleus was measured by ImageJ. B. Western blot analysis showing the knockdown efficiency for macroH2A1.1 and macroH2A1.2. Bands were quantitated from four independent experiments using Syngene software. B. Levels of macroH2A1 was normalized to histone H3 and then normalized to T=0 The paired student t test was used to determine statistical significance. Errors bars represent +/- SEM, *, P<0.05; **P<0.005; *** P<0.005

**Supplementary Figure 3.**
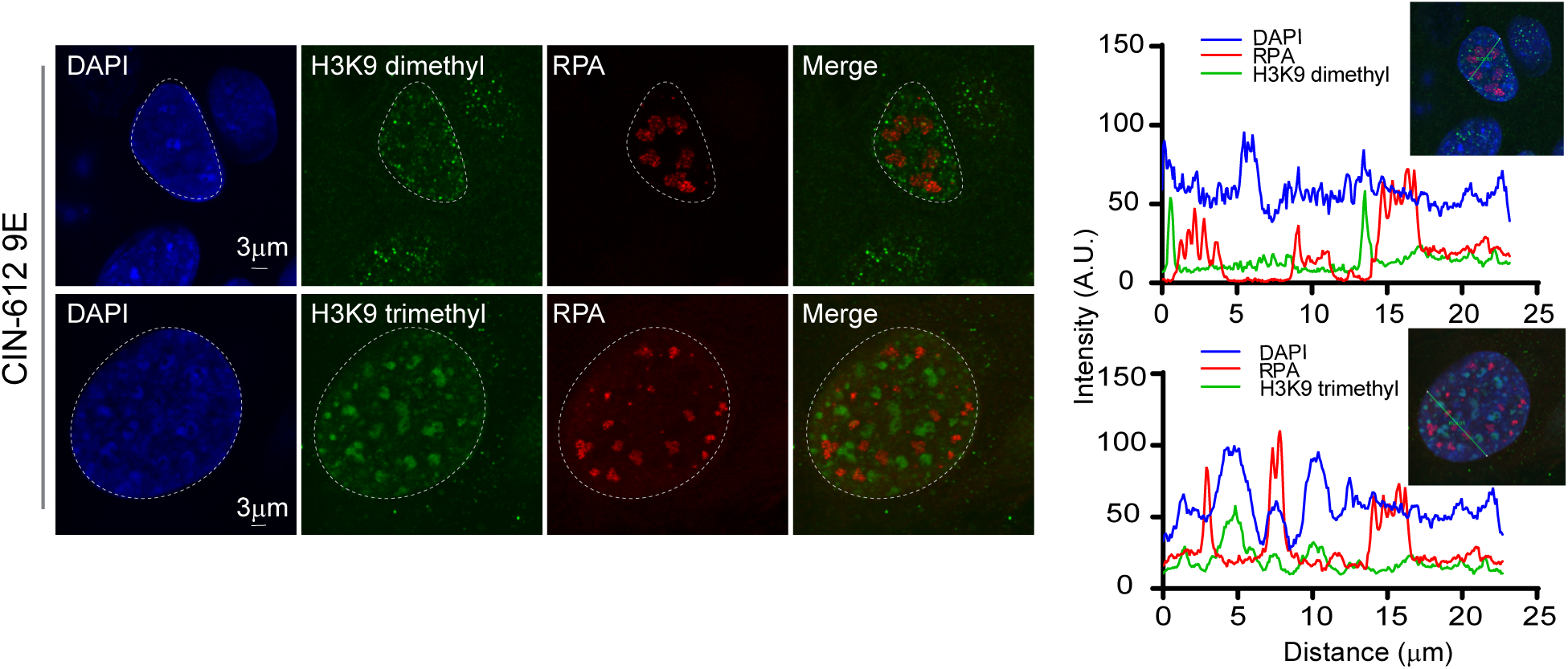
Histone modifications associated with repressed chromatin are not enriched at viral replication foci in differentiated 9E cells. Differentiated 9E cells were immunostained with antibodies against histone H3K9 dimethyl or trimethyl (green) and RPA (red). The nuclei were stained with DAPI (blue). For H3K9 dimethyl, a total of 38 cells (N= 38) and 144 RPA positive foci were counted and for H3K9 trimethyl a total of 39 cells (N=39) and 107 RPA positive foci were counted in two independent experiments.

**Supplementary Figure 4.**
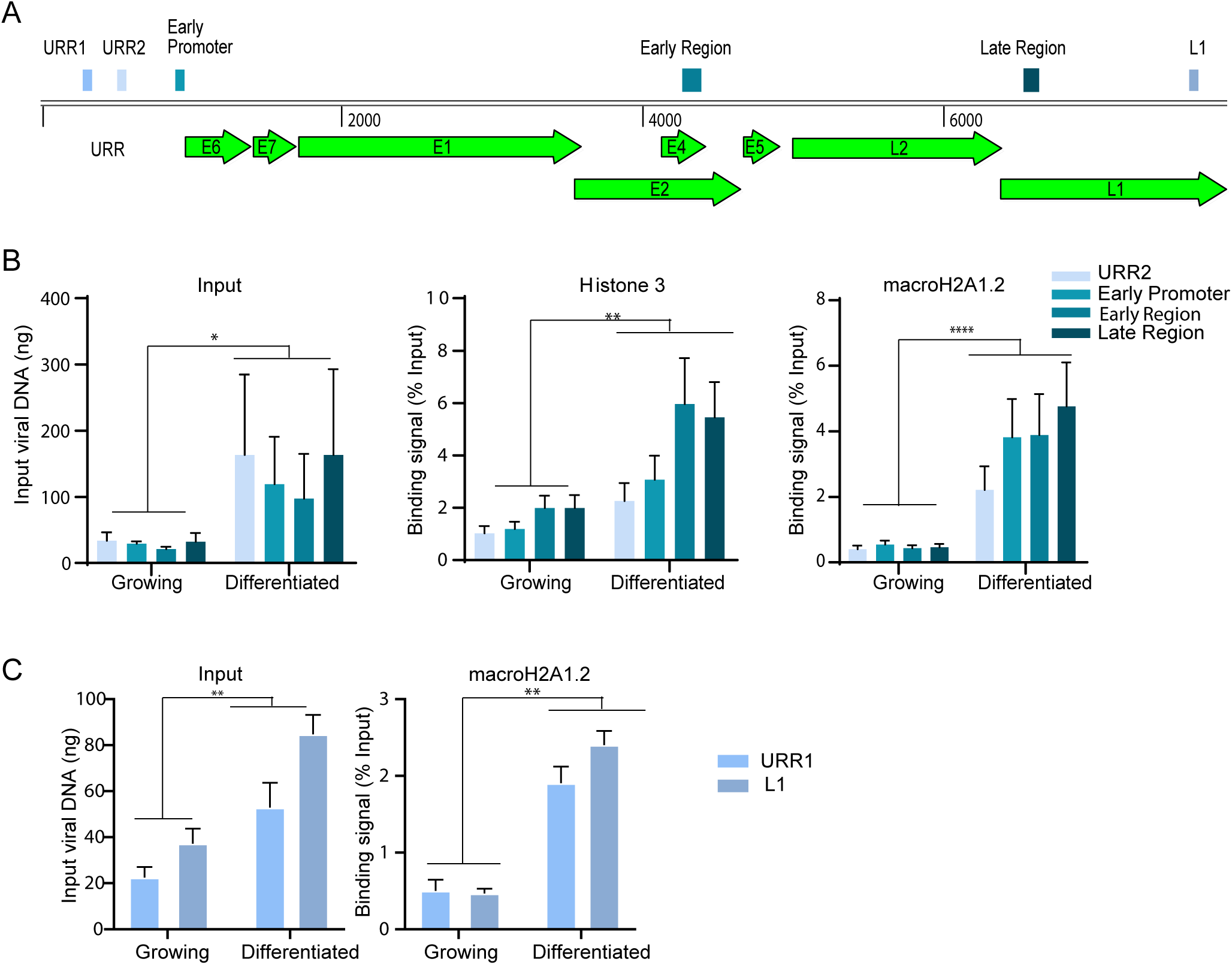
macroH2A1.2 binding to HPV chromatin increased upon differentiation. Chromatin immunoprecipitation (ChIP) was carried out with chromatin isolated from growing and differentiated 9E cells using rabbit IgG and macroH2A1.2 antibodies. **A.** Schematic of HPV31 genome showing primer positions for upstream regulatory region, early promoter, early region and late region**. B.** ChIP signals for histone H3 and macroH2A1.2 were expressed as a percentage of immunoprecipitated viral DNA relative to the total amount of input chromatin. Background signal (measured by immunoprecipitating viral DNA with IgG antibody) was subtracted from the corresponding ChIP signals. Binding levels were averaged from three independent experiments. Error bars represent +/- SEM and statistical significance was calculated using a paired student t-test. **C**. ChIP signals were expressed as a percentage of immunoprecipitated viral DNA relative to the total amount of input chromatin. Background signal (measured by immunoprecipitating viral DNA with IgG antibody) was subtracted from the corresponding ChIP signals.URR1 and L1 regions were selected from ChIP seq peaks and ChIP was carried out independent of ChIP experiment shown in B. Binding levels were averaged from two independent experiments. Error bars represent +/- SD and statistical significance was calculated using a paired student t-test. **\***P<0.05; **P<0.005; *** P<0.005

**Supplementary Figure 5:**
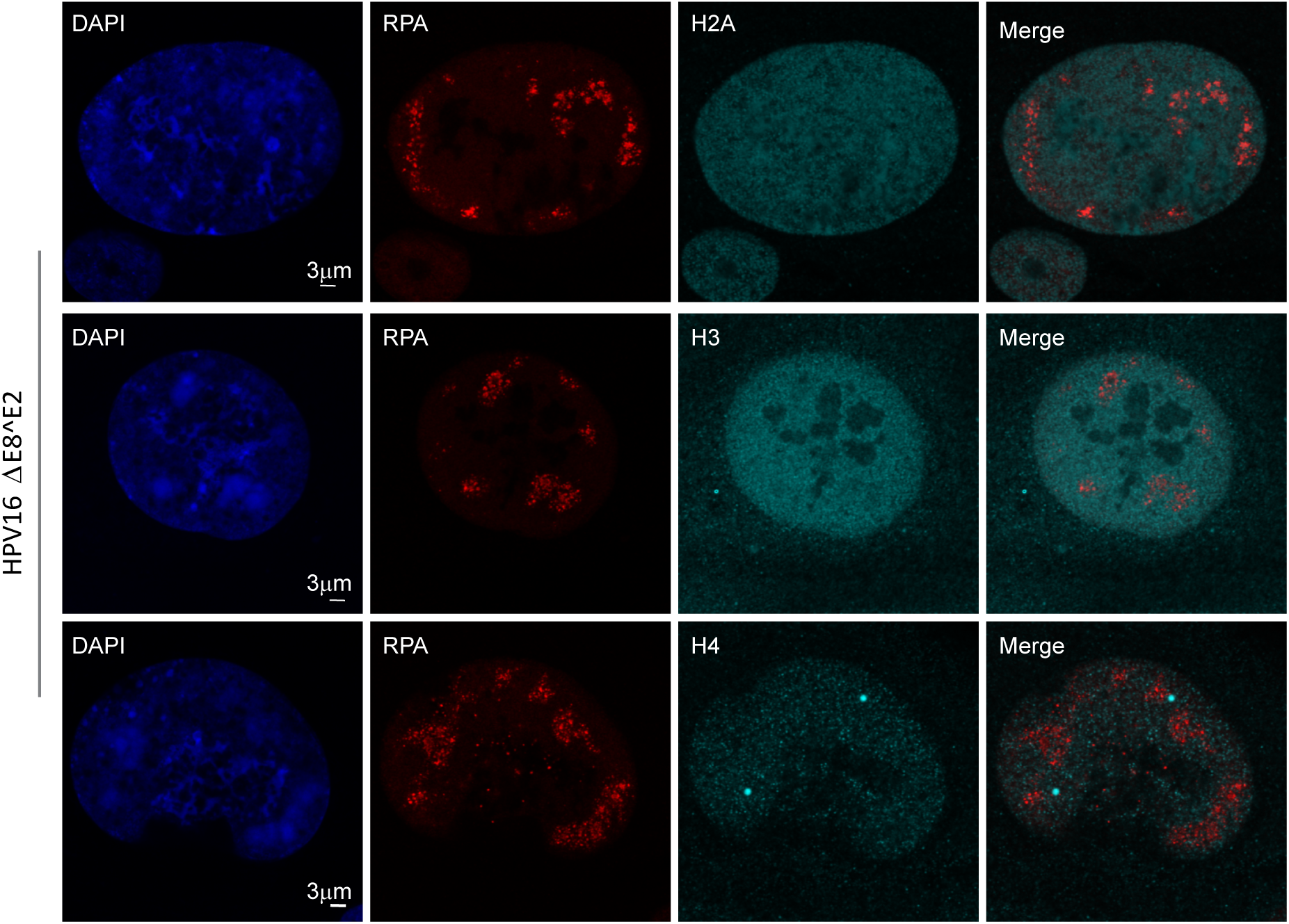
Canonical histones are neither depleted nor enriched in HPV16 ΔE8^E2 replication foci. HPV16 ΔE8^E2 genome containing cells were immunostained with antibodies against histones H2A, H3 and H4 (cyan), macroH2A1.2 (green) and RPA (red). Nuclei were stained with DAPI (blue) and the distribution of histone H2A (N=35 cells, N=162 RPA positive foci), histone H3 (N=43 cells, N=171 RPA positive foci) and histone H4 (N=34 cells, N=162 RPA positive foci) was analyzed. Cells and foci were scored from two independent experiments.

**Supplementary Figure 6:**
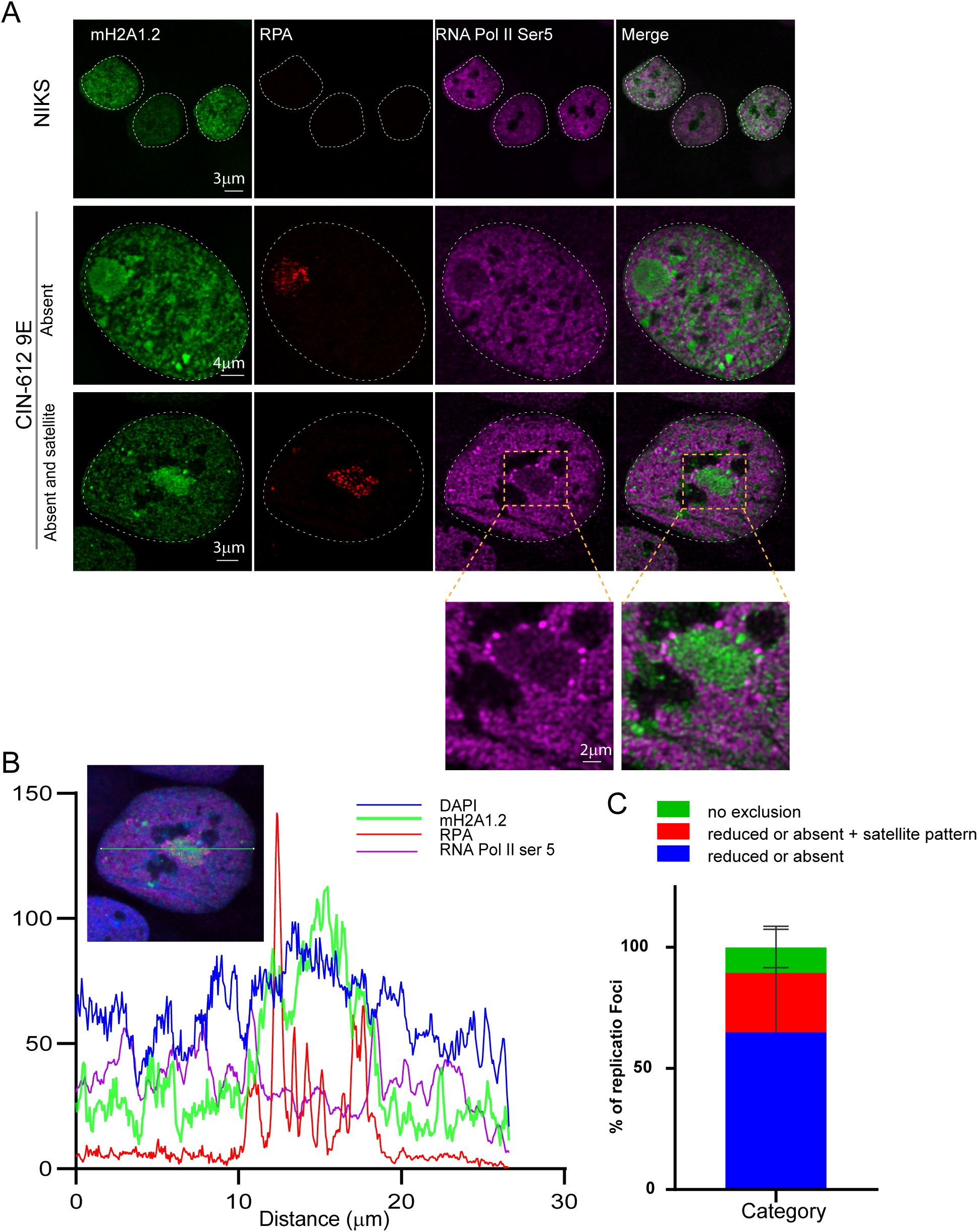
RNA Pol II Ser 5 is excluded from the replication foci in differentiated 9E cells. **A**. Differentiated NIKS or 9E cells were immunostained with antibodies against macroH2A1.2 (green), RNA Pol II Ser 5 (purple) and RPA (red). A white dotted line outlines the nuclei. In differentiated CIN-612 9E cells, (N=71), 233 foci were counted with RPA used as a marker for viral replication foci. In differentiated NIKS cells (N=34) cells were negative for foci. Data are from two independent experiments. **B**. Fluorescence intensity scan obtained by drawing a line through a nucleus shown in left panel. **C.** Distribution of RNA Pol II Ser5 at replication foci by visual counting (N=71, total of 233 foci counted) in two independent experiments.

**Supplementary Table 1:**
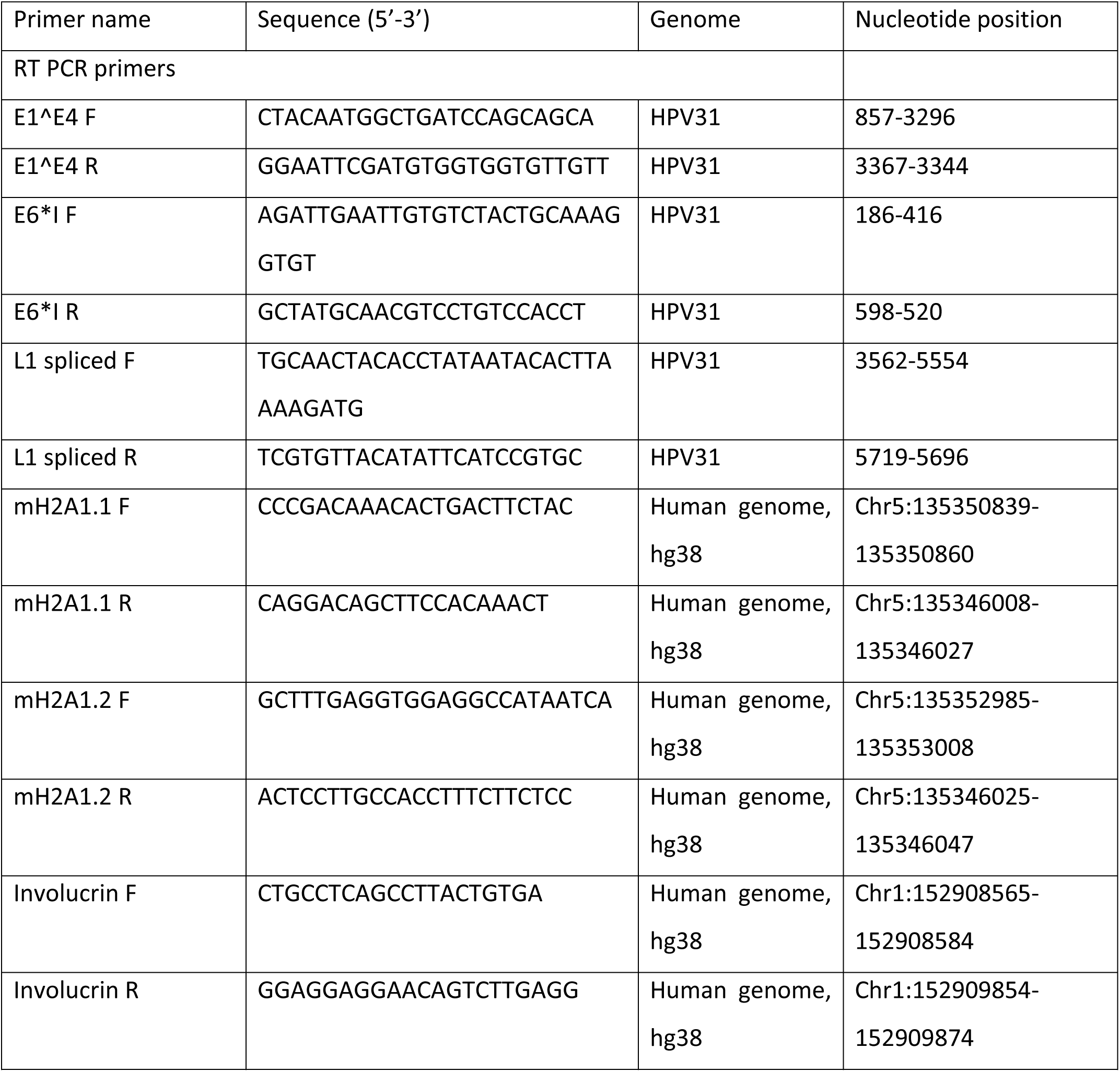
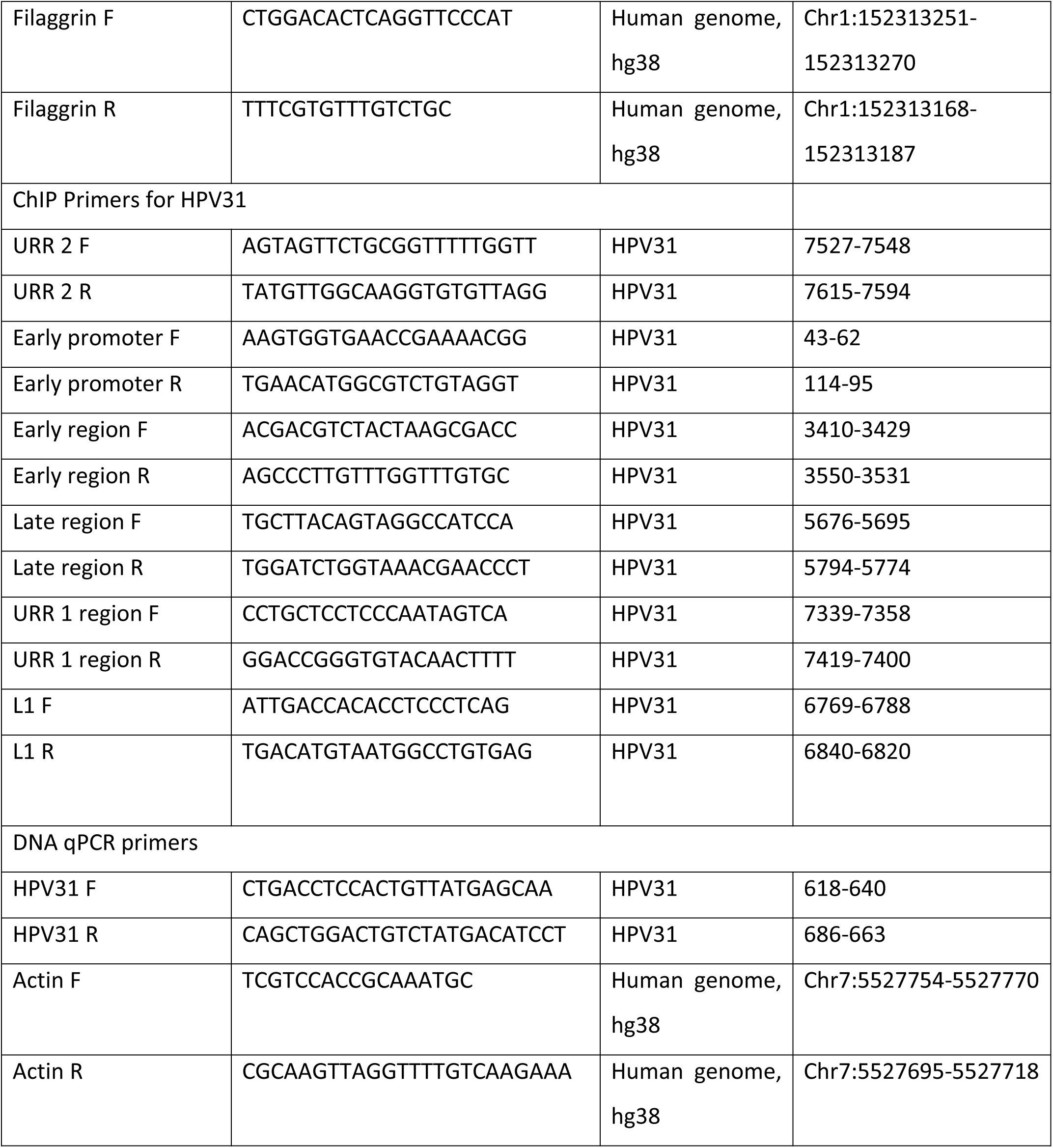

